# Predicted Effect of Circadian Regulation of Renal Transporters in a Hypertensive Rat

**DOI:** 10.1101/2024.12.11.628058

**Authors:** Kaixin Zheng, Anita T. Layton

**Author notes:** **Corresponding author:** Anita Layton Department of Applied Mathematics University of Waterloo Waterloo, Ontario Canada, N2L 3G1.

## Abstract

The kidney regulates extracellular fluid and electrolyte homeostasis. Its function is modulated by the renin-angiotensin-aldosterone system (RAAS). Overactivation of the RAAS, such as in chronic Ang II infusion, promotes vasoconstriction, anti-natriuresis, and hypertension. Kidney function is also modulated by circadian clocks. Indeed, glomerular filtration rate, filtered electrolyte loads, urine volume, and urinary excretion all exhibit notable diurnal rhythms, which reflect, in part, the regulation of renal transporter proteins by circadian clock genes. Key renal transporters that are regulated by clock gene proteins include sodium-hydrogen exchanger 3 (NHE3), sodium-glucose cotransporter 1 (SGLT1), Na^+^-K^+^-2Cl^−^ cotransporter (NKCC2), Na^+^-Cl^−^ cotransporter (NCC), and epithelial sodium channel (ENaC), which are targeted by diuretics used to treat hypertension. The objective of the present study was to assess the effect of administration time on the natriuretic and diuretic effects of loop, thiazide, and K^+^-sparing diuretics, which are common treatments for hypertension, and how those effects differ between a normotensive and hypertensive kidney. Loop diuretics inhibit NKCC2 on the apical membrane of the thick ascending limb; notably, In Ang II-induced hypertension, NKCC2 is differentially regulated along the medullary versus cortical thick ascending limb. Thiazide diuretics inhibit NCC on the distal convoluted tubule, and K^+^-sparing diuretics inhibit ENaC on the connecting tubule and collecting duct. We simulated Na^+^ transporter inhibition using computational models of kidney function at different times of day. The simulation results predicted qualitatively similar diurnal oscillations in segmental transport in the normotensive and hypertensive kidneys, and highlighted significant time-of-day differences in the natriuresis, diuresis, and kaliuresis responses.

**NEW & NOTEWORTHY:** Chronic infusion of angiotensin II promotes vasoconstriction, salt retention, and hypertension. Blood pressure and kidney function exhibit circadian rhythms. Given the diurnal variations in the expression levels of key renal electrolyte transporters, how do the natriuretic and diuretic effects of diuretics, a common treatment for hypertension that targets renal transporters, vary during the day? To answer this question, we simulated Na^+^ transporter inhibition using computational models of kidney function at different times of day.

## INTRODUCTION

The kidney is a major regulator of extracellular fluid and whole-body electrolyte homeostasis. In particular, the kidney adjusts the fraction of Na^+^ in its glomerular filtrate that it reabsorbs, to match the amount of Na^+^ excreted in the urine with Na^+^ intake. Given that the kidneys receive 20-25% of the cardiac output (1), glomerular filtration rate (GFR) and filtered Na^+^ load are high, and to achieve Na^+^ balance, only about 1% of the filtered Na^+^ is excreted in the urine (in a normal kidney) (2). This necessitates a highly precise adaptation of the renal transport system. Indeed, almost all nephron segments participate in the reabsorption of filtered Na^+^. Along the proximal tubule, the Na^+^/H^+^ exchanger 3 (NHE3) mediates the reabsorption of a large fraction of the filtered Na^+^ (about 50%-70%, more in male rodents) (3). The NHE3 also plays a key role in the pressure natriuresis response, whereby an increase in blood pressure leads to an increase in Na^+^ excretion (4). The thick ascending limb is another major Na^+^-reabsorbing segment, where the Na^+^-K^+^-2Cl^-^ cotransporter 2 (NKCC2) on the apical membrane is responsible for 25%-40% of the Na^+^ transport (more in females) (5,6). The importance of NKCC2 for Na^+^ balance can be seen in the powerful antihypertensive effect of loop diuretics, which inhibit NKCC2. Among the downstream segments, Na^+^-Cl^-^ cotransport (NCC) on the apical membrane of the distal convoluted tubule mediates Na^+^ uptake, as does the epithelial Na^+^ channel (ENaC) along the principal cells of the connecting tubule and collecting duct (7). These segments are responsible for “fine-tuning” the final urinary excretion, and the importance of NCC and ENaC in Na^+^ balance is evinced by the extensive use of thiazide diuretics and K^+^-sparing diuretics, which target NCC and ENaC, respectively, in treatment of hypertension.

The maintenance of fluid and electrolyte homeostasis by the kidney is modulated by the renin-angiotensin-aldosterone system (RAAS). An overactive RAAS may lead to Na^+^ retention, K^+^ loss, and an increase in blood pressure. In particular, angiotensin II (Ang II) regulates renal Na^+^ transport at a molecular level, and chronic infusion of Ang II induces vasoconstriction and anti-natriuresis (8,9). Salt retention is a consequence of the Ang II-induced changes in key renal electrolyte transporters (10–12): downregulation of NHE3 and NKCC2 in the proximal tubule and medullary thick ascending limb, upregulation of NKCC2 in the cortical thick ascending limb, NCC, and ENaC. These changes result in a downstream shift in Na^+^ transport capacity of the nephron, Na^+^ and fluid imbalance, and hypertension (10,13,14).

Describing a primary function of the kidney as to maintain the “homeostasis” of fluid and electrolyte may give the impression that with sufficient adjustments and feedback control, a steady state or equilibrium can be achieved. However, this picture is incomplete: the kidney, like most physiological systems, exhibits circadian rhythms with a 24-h period. In the mammalian kidney, the circadian rhythms are co-mediated by the central clock, which resides in the suprachiasmatic nucleus (SCN) of the hypothalamus, and by the peripheral clocks within the renal cells that can oscillate independently of the SCN (15). Both the central and peripheral clocks ‘tick” as a result of the interactions among a network of core clock genes (16–18). Briefly, the basic helix-loop-helix ARNT like 1 (Bmal1) and Clock dimerize to induce the transcription of Period (Per) and Cryptochrome (Cry) genes. The proteins PER and CRY then heterodimerize to act on the protein complex CLOCK-BMAL1, thereby inhibiting their own transcription.

This feedback loop of the core clock genes results in oscillations in protein levels, which drive circadian rhythms observed in kidney function, including the marked reduction in the volume of urine excreted during the night (in diurnal animals) compared to the volume excreted during the day (19,20). Similar oscillations are observed in the urinary excretion of electrolytes, renal plasma flow. and glomerular filtration rate. The circadian rhythms of key kidney function arise in part from the regulation by clock proteins of renal transporter genes, including those of NHE3, Na^+^-glucose cotransporter 1 (SGLT1), NKCC2, NCC, and ENaC (19,20).

As previously noted, NKCC2, NCC, and ENaC are the targets of loop, thiazide, and K^+^-sparing diuretics, common medications for hypertension (21). Given the diurnal variations in the expression levels of these transporters, how do the natriuretic and diuretic effects of these diuretics vary during the day? To answer this question, we simulated Na^+^ transporter inhibition using our recently published computational models of water and electrolyte transport along the nephrons of a male kidney that represent the regulation of transporter activities by circadian clock proteins.

## MATERIALS & METHODS

### Modeling the circadian clock network

We aim to simulate the modulation of key renal transporter activities in a male rat by the circadian clock. We first develop a mathematical model of the core clock in the male rat kidney. The clock model comprises a number of transcription factors that regulate gene expression: the period homologs Per1 and Per2, the cryptochromes homologs Cry1 and Cry2, Rev-Erb and RAR-related orphan receptor (Ror), brain and muscle ARNT-Like 1 (Bmal1), and circadian locomotor output cycles kaput (CLOCK). A schematic diagram of the clock network is shown in Fig. 1. Equations for the clock model are given in Eqs. A1-A25. Model parameters were obtained by fitting predicted profiles for mRNA expression levels of core clock genes (Per1, Per2, Cry1, Cry2, Rev-Erb, Ror, Bmal1) with their corresponding measured data reported in Ref. (22) and database (23), obtained for the dark-dark cycles.

**Figure 1.**
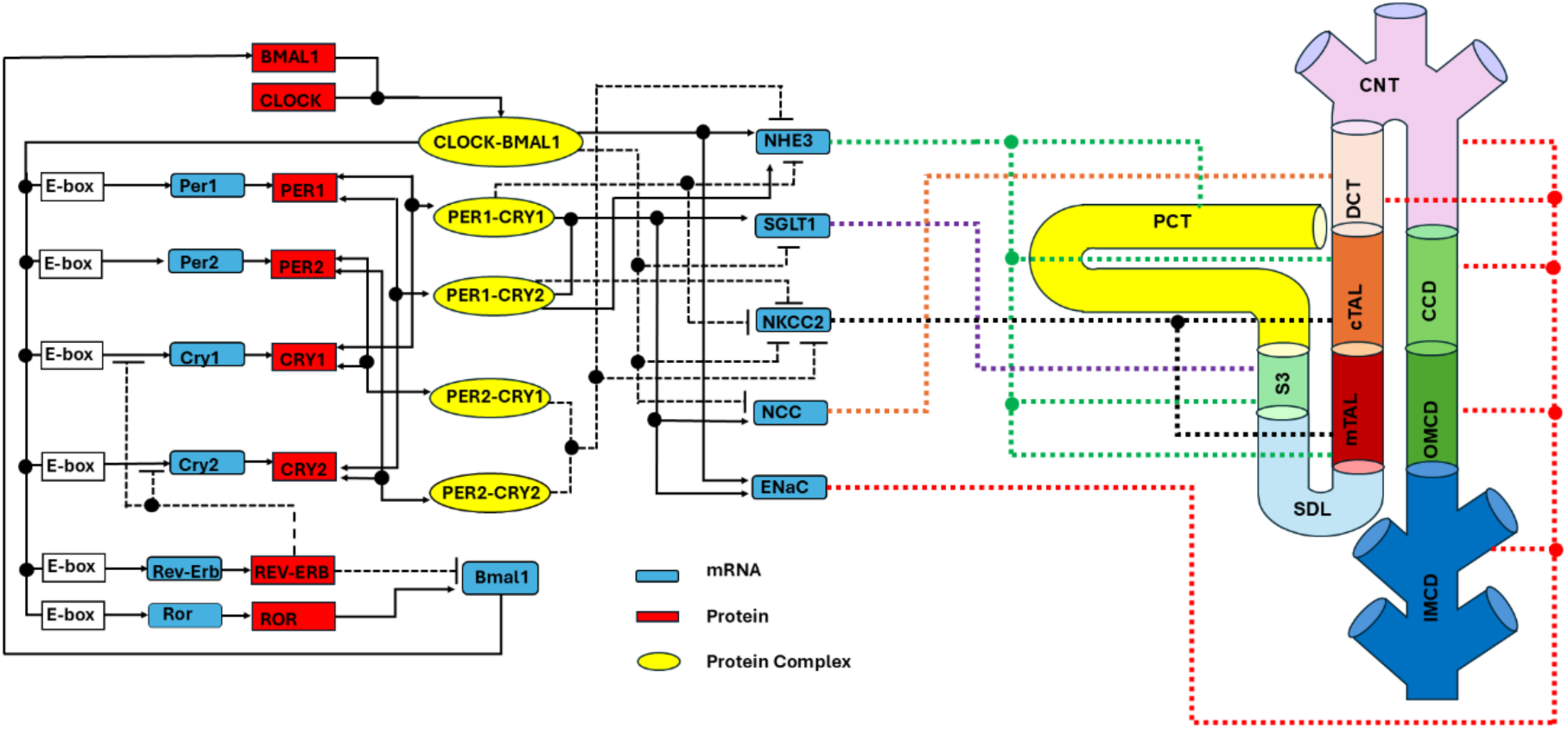
Schematic network of our mathematical model, which describes the interactions of core clock components with key renal transporter genes. In the core clock network, mRNAs are denoted by blue boxes, proteins by red boxes, and protein complexes by yellow ovals. Solid lines represent the transactivation and dotted lines represent inhibition. The model nephron includes the proximal convoluted tubule (PCT), S3 segment, medullary thick ascending limb (mTAL), cortical thick ascending limb (cTAL), distal convoluted tubule (DCT), connecting tubule (CNT), cortical collecting duct (CCD), outer medullary collecting duct (OMCD), and inner medullary collecting duct (IMCD). Renal transporters that are regulated by the circadian clock include NHE3, SGLT1, NKCC2, NCC, and ENaC.

### Modeling renal epithelial transport and its circadian regulation

The model represents a superficial nephron of a male rat kidney. As depicted in Fig. 1, the model nephron is divided into 10 functionally distinct segments: proximal convoluted tubule (PCT), proximal straight tubule (S3), short descending limb (SDL), medullary thick ascending limb (mTAL), cortical thick ascending limb (cTAL), distal convoluted tubule (DCT), connecting tubule (CNT), cortical collecting duct (CCD), outer medullary collecting duct (OMCD), and inner medullary collecting duct (IMCD). Each segment is modeled with a series of computational cells assigned with specific physical dimensions, transporter profile, and membrane permeability. The model accounts for the following 15 solutes: Na^+^, K^+^, Cl^−^, HCO_3_^−^, H_2_CO_3_, CO_2_, NH_3_, NH_4_^+^, HPO_4_^2−^, H_2_PO_4_^−^, H^+^, HCO_2_^−^, H_2_CO_2_, urea, and glucose. In each computational cell, steady-state luminal, cellular, and paracellular concentrations and fluxes are calculated based on water conservation, nonreacting solute conservation, and pH conservation. Model equations can be found in Ref. (24).

We represent the circadian regulation of key Na^+^ transporters. Specifically, NHE3 expressed on the PCT, S3, mTAL, and cTAL, sodium-glucose cotransporter 1 (SGLT1) on the S3, NKCC2 on the mTAL and cTAL, NCC on the DCT, and ENaC along the DCT, CNT, and CD.

Rohman et al. (25) experimentally verified that that NHE3 in the rat kidney is transactivated by CLOCK-BMAL1, and that Cry1 and Per2 inhibit the CLOCK-BMAL1-mediated transactivation of the NHE3 promoter, with the inhibition much stronger for Cry1. Solocinski et al. (26) demonstrated that Per1 enhances NHE3 expression. Therefore, we assume that NHE3 is activated by CLOCK-BMAL1 and PER1-CRY2, but inhibited by PER1-CRY1, PER2-CRY1 and PER2-CRY2.

Crislip et al. (27) reported a significant increase in NKCC2 mRNA expression in Bmal1 knockout male mice compared to wild-type. Thus, we assume that NKCC2 is inhibited by CLOCK-BMAL1. Given that NKCC2 mRNA exhibits sustained oscillations in Bmal1 knockout mice (27), we assume additionally that NKCC2 is also inhibited by PER1-CRY1, PER1-CRY2, PER2-CRY1, and PER2-CRY2. Supporting experimental evidence is limited and indirect: estrogen-related receptor beta (ERRβ), which displays a circadian rhythm in the kidney with peak expression at ZT4, is known to positively regulate NKCC2 expression in male mice (28), and estrogen receptor beta (ERβ) is downregulated by the PER-CRY complexes (29).

SGLT1 is assumed to be activated by PER1-CRY1 and PER1-CRY2 but inhibited by CLOCK-BMAL1. This assumption is supported by evidence that Per1 knockout decreases SGLT1 expression in male mice (26), and that BMAL1 inhibits SGLT1 in Caco-2 cells via activation of paired-homeodomain transcription factor 4 (PAX4) (30). Additionally, Stevenson et al. (31) demonstrated that activation of Bmal1 decreases renal SGLT1 levels in an SHP-dependent manner, whereas Clock-mutant and Bmal1-knockout mice show elevated SGLT1 expression, supporting the idea that CLOCK-BMAL1 inhibits SGLT1.

In both male and female mouse kidneys ,NCC expression is positively regulated by WNK4 (32), the expression of which increases significantly in male Bmal1-knockout mice compared to wild-type (27). Also, Per1 blockade inhibits NCC activity in male mice (33). Thus, in the model NCC expression is suppressed by CLOCK-BMAL1 and activated by PER1-CRY1 and PER1-CRY2.

Based on findings in mice (34), ENaC is assumed to be activated by CLOCK-BMAL1 and also by PER1-CRY1 and PER1-CRY2, which colocalizes with CLOCK-BMAL1.

Transporter activities are assumed proportional to the corresponding mRNA expression levels, and thus fluctuates with core clock protein levels. The links are described in Eqs. A8-A12 and summarized in Fig. 1. Model parameters were obtained by fitting predicted profiles for the transporters (NHE3, SGLT1, NKCC2, NCC, ENaC) with data reported in Refs. (27,35,36).

### Modeling the circulation regulation of renal hemodynamics

As was done in Ref. (36), the model represents the single-nephron GFR (SNGFR) of a wild-type male rat in a light-dark cycle, denoted SNGFR_!"_, as a sinusoidal function of the Zeitgeber Time (ZT), denoted t, with a peak at ZT16:

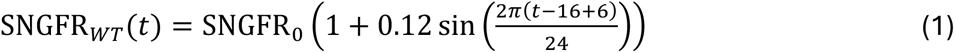

ZT=0 (lights on) marks the start of the rest phase for nocturnal animals, whereas ZT12 (lights oX) denotes the start of the active phase. The baseline SNGFR_0_is taken to be 32 nl/min for the superficial nephron of a male rat.

GFR in Bmal1 knockout mice exhibits an ultradian rhythm different from wile-type, with two peaks at ZT4 and ZT16 ; the 24-hour cumulative GFR are similar in the two genotypes (37). Based on these findings, we model SNGFR in Bmal1-knockout rat, denoted SNGFR_./_, as:

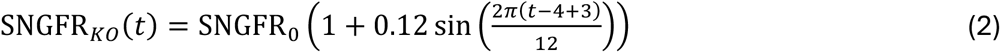

### Modeling the effects of hypertension on renal epithelial transport

We investigate the effect on kidney function in a male rat with hypertension induced by 14-day infusion of ANG II. Following the approach in Ref. (38), we incorporate hypertension-induced changes into the renal epithelial transport model described above. These changes include changes in transport activities, membrane permeabilities, and interstitial fluid composition. Among the transporters of which the circadian regulation is represented in the present model, NHE3 and NKCC2 along the mTAL are downregulated in hypertension; NKCC2 along the cTAL, NCC, and ENaC are upregulated, whereas SGLT1 remains unchanged. The links between clock proteins and transporter activities are assumed the same in normotension and hypertension, as is the SNGFR profile.

### Modeling the effects of diuretics on renal epithelial transport

We consider the effects of loop diuretics, thiazide diuretics, and K^+^-sparing diuretics on renal transport and excretions. Loop diuretics inhibit NKCC2, and its administration is assumed to significantly impair the model kidney’s ability to generate an axial osmolality gradient. Thus, we lower the interstitial fluid concentrations of selected solutes (39), but keep the interstitial urea concentration at the papillary at the baseline hypertension value (which is assumed to be lower than in normotension) (38). SNGFR is assumed to retain its baseline profile.

Thiazide diuretics inhibit NCC. We simulate a dosage of thiazide diuretics that induces 100% inhibition of NCC. All other model parameters remain at baseline values. Interstitial concentration profiles are assumed unchanged.

K^+^-sparing diuretics inhibit ENaC. We simulate a dosage of K^+^-sparing diuretics that induces 100% inhibition of ENaC. Interstitial concentration profiles are again assumed unchanged.

## RESULTS

### Predicted diurnal oscillations of clock gene and transporter expression levels

Using the baseline model parameters (see Appendix), the circadian clock network model predicts that the expression levels of all core clock components exhibit limit-cycle oscillations with a 24-h period. Time-profiles of core clock components, together with the experimental data (22), are shown in Fig. 2. Model parameters were chosen to ensure good agreement between the predicted profiles and data.

**Figure 2.**
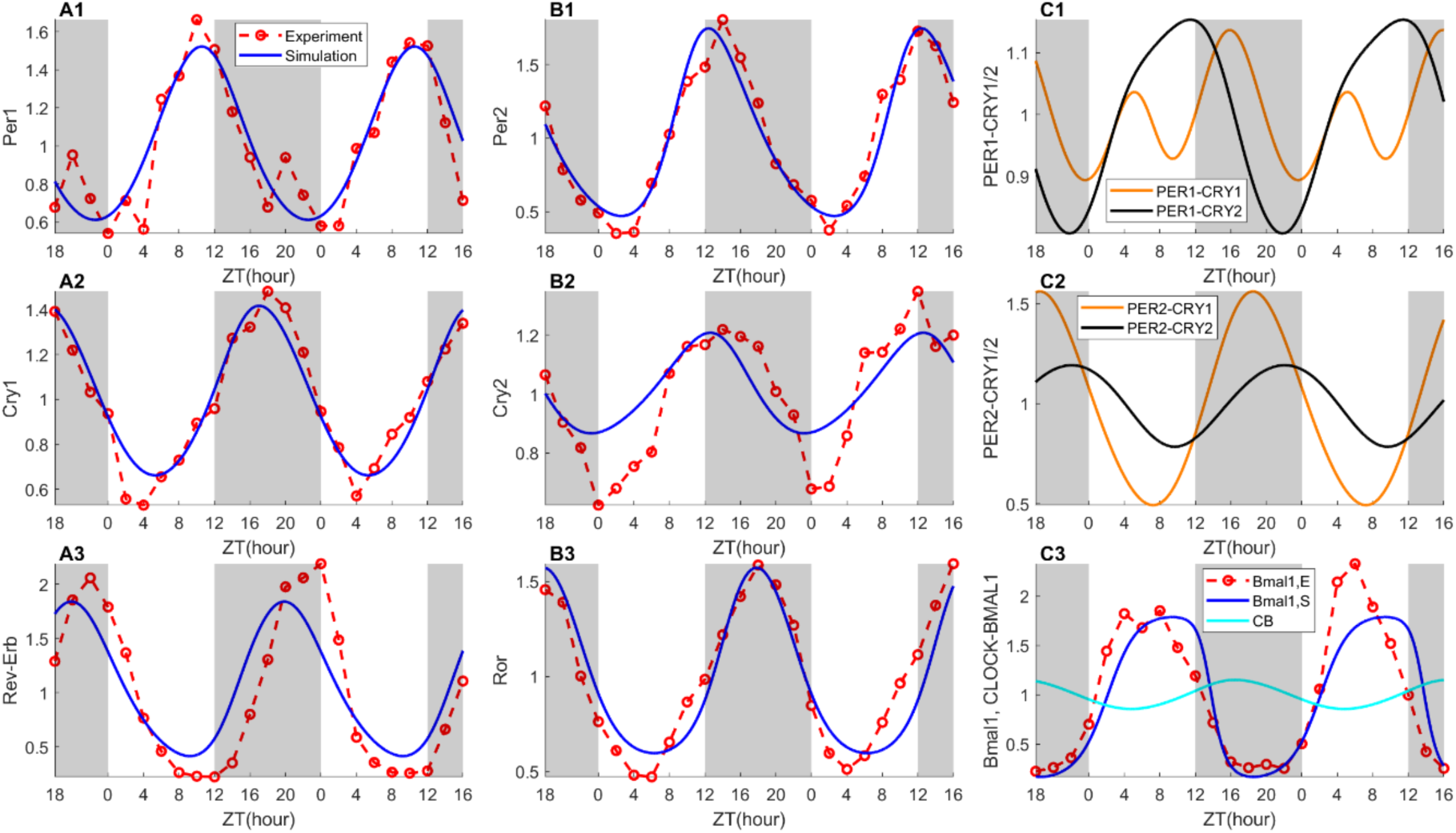
Comparison of simulated circadian gene and protein profiles with data from 46-hour experiments starting at ZT18 (22,23). ***A1***–***A3***, ***B1***–***B3***, simulated mRNA expression levels (Simulation) compared with experimental data (Experiment), normalized by mean experimental values; ***C1***, ***C2***, simulated concentrations of PER1–CRY1, PER1–CRY2, PER2– CRY1, and PER2–CRY2 protein complexes, normalized by their respective mean simulated values; ***C3***, simulated BMAL1 mRNA expression levels (Bmal1, S), experimental BMAL1 mRNA levels (Bmal1, E), and normalized CLOCK–BMAL1 complex concentrations (CB). Gray shading and white regions correspond to dark and light phase, respectively.

Oscillations in the core clock components drive oscillations in the expression levels of NHE3, SGLT1, NKCC2, NCC, and ENaC. We identify parameters that describe the regulation of NHE3 expression by CLOCK-BMAL1 and the PER-CRY complexes that the predicted NHE3 time profiles agree with the normotensive male mouse data in the database (23) from the mice experiment (22) in constant dark environment. As shown in Fig. 3A, the predicted NHE3 expression profile exhibits good agreement with experimental data. Similarly, model parameters that link the expression levels of SGLT1, NCC, and ENaC to core clock proteins were fitted, so that the predicted transporter expression levels approximate rat data obtained in light-dark cycles (35). Results are shown in Fig. 3B, 3C, and 3D.

**Figure 3.**
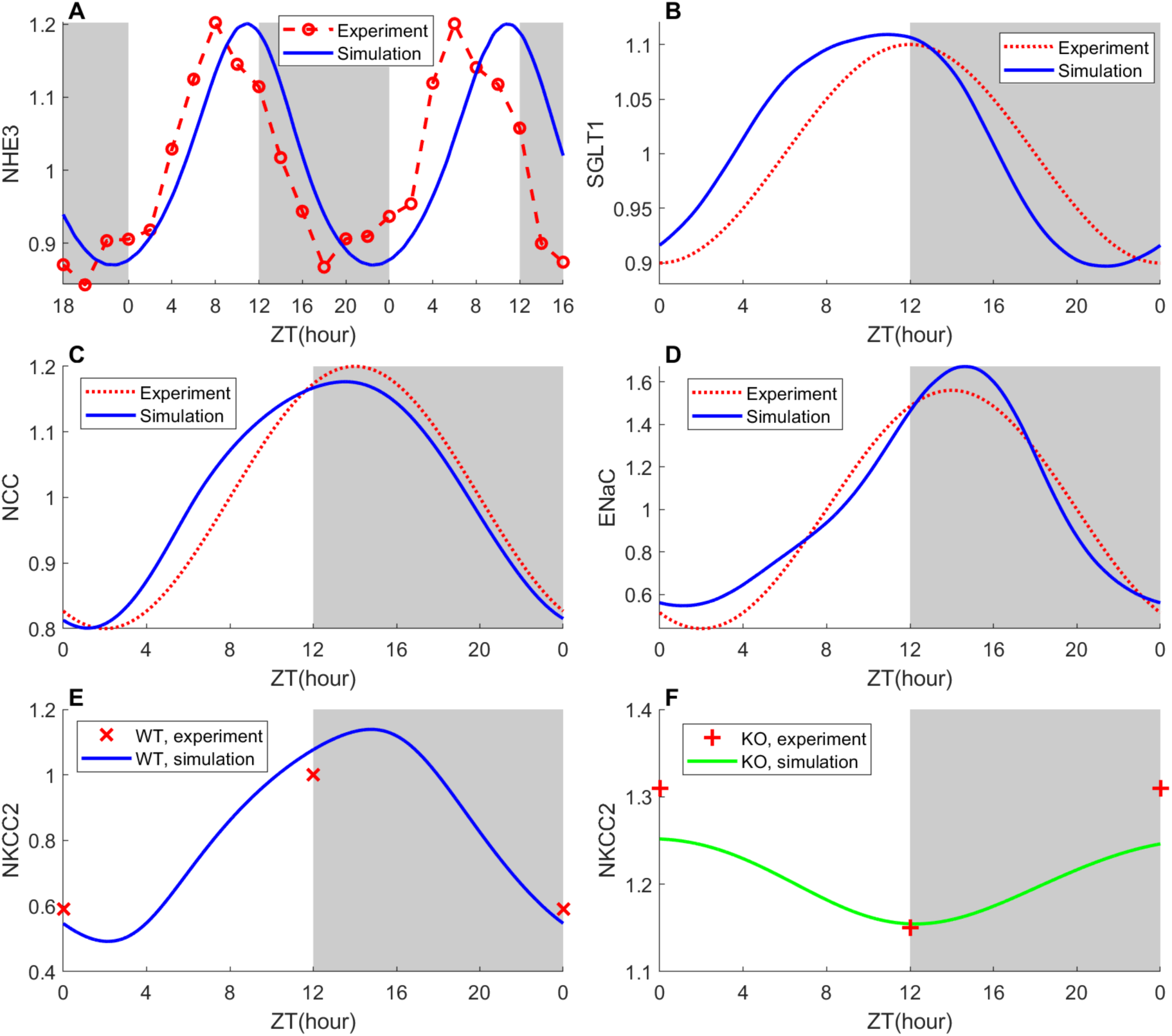
Comparison of simulated transporter expression with corresponding experimental data. ***A***-***D***, Transporter expression simulation (Simulation) and experimental profiles (Experiment) of NHE3, SGLT1, NCC, ENaC respectively. ***E***, Simulation of NKCC2 expression in wild-type (WT, simulation) and corresponding experimental data at noon and midnight (WT, experiment). ***F***, Simulation of NKCC2 expression after Bmal1 knockout (KO, simulation) with corresponding experimental data at noon and midnight (KO, experiment). Values are normalized by mean expression level in wild type. Gray shading and white regions correspond to the dark and light phases, respectively.

Like NHE3, NKCC2 expression is also regulated by CLOCK-BMAL1 and the PER-CRY complexes. We fit those parameters against data measured in wild-type and Bmal1-knockout rats (27). To simulate the renal peripheral clock in Bmal1-knockout rats, we reduce the maximal transcription rate for Bmal1 by 40%. We then compare the predicted NKCC2 expression levels in these two groups at noon and midnight with values reported in Ref. (27). Results are shown in Fig. 3E and 3F. Using these parameters, the predicted NKCC2 levels for both wild-type and Bmal1-knockout align with experimental data obtained at noon and midnight. The suppression of Bmal1 transcription is predicted to cause a significant phase shift in NKCC2 oscillations: the peak time occurs at ZT15 in wild-type, but shifts to ZT0 in Bmal1-knockout. Additionally, Bmal1 knockout substantially increases average NKCC2 expression (46%), whereas the oscillation amplitude decreases (85%).

The amplitudes and peaks (normalized by means) of the predicted core clock gene and transporter mRNA time profiles for the wild-type model are summarized in Table 1.

**Table 1.**
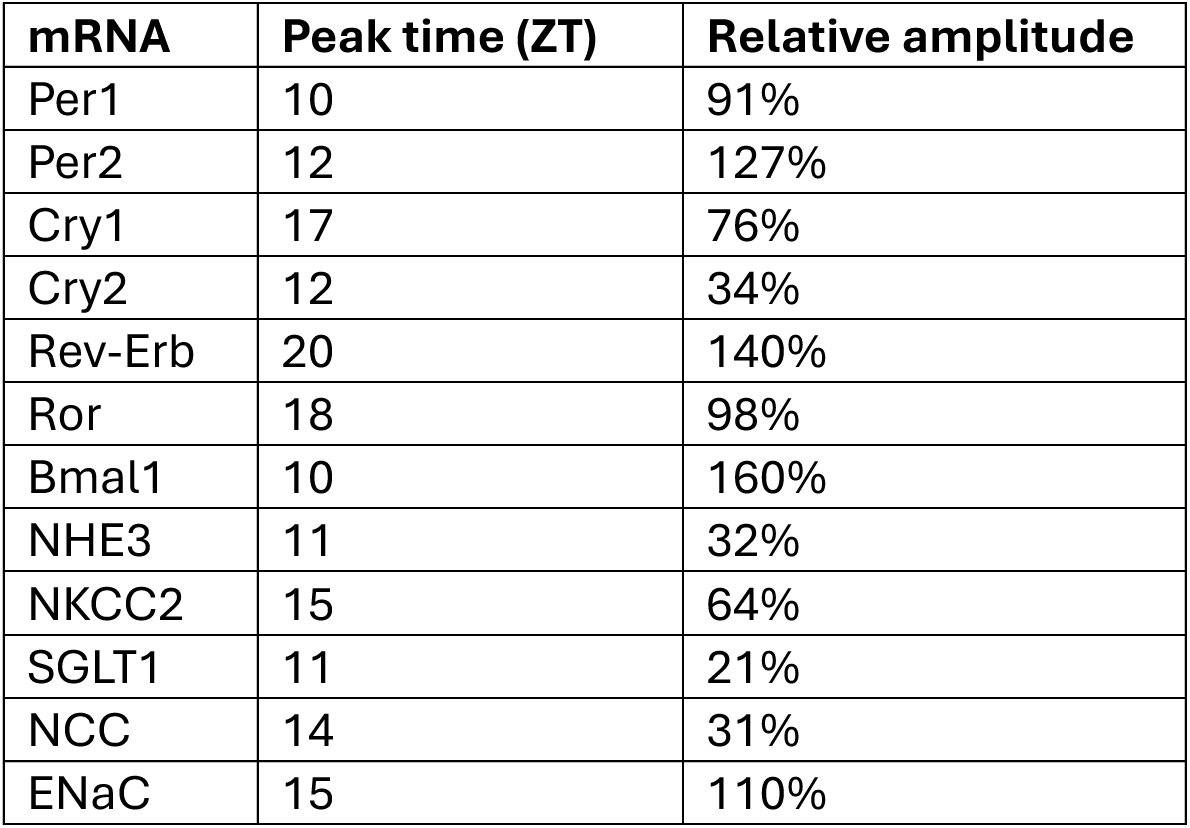
Peak time and relative amplitude of predicted mRNA oscillations in the kidneys of wild-type normotensive male rats. Relative amplitude is calculated as (peak - trough)/mean.

| **mRNA** | **Peak time (ZT)** | **Relative amplitude** |
| --- | --- | --- |
| Per1 | 10 | 91% |
| Per2 | 12 | 127% |
| Cry1 | 17 | 76% |
| Cry2 | 12 | 34% |
| Rev-Erb | 20 | 140% |
| Ror | 18 | 98% |
| Bmal1 | 10 | 160% |
| NHE3 | 11 | 32% |
| NKCC2 | 15 | 64% |
| SGLT1 | 11 | 21% |
| NCC | 14 | 31% |
| ENaC | 15 | 110% |

### Predicting segmental Na^+^ transport and urinary excretion in Bmal1-knockout rats

We then investigate how Bmal1 knockout affects nephron function at different times of day. To that end, we incorporate the predicted time profiles of NHE3, SGLT1, NKCC2, NCC, and ENaC (Fig. 3), together with the time-varying SNGFR (Eqs. 1 and 2), into the nephron transport model, and we conduct simulations to predict water and solute transport along individual nephron segments and urine excretions, at different ZT. As noted above, we simulate conditional Bmal1 knockout by reducing the maximal transcription rate of Bmal1 by 40%. SNGFR and the predicted transporter profiles are shown in Fig. 4. Segmental Na^+^ reabsorption and Na^+^ excretion are summarized in Fig. 5.

**Figure 4.**
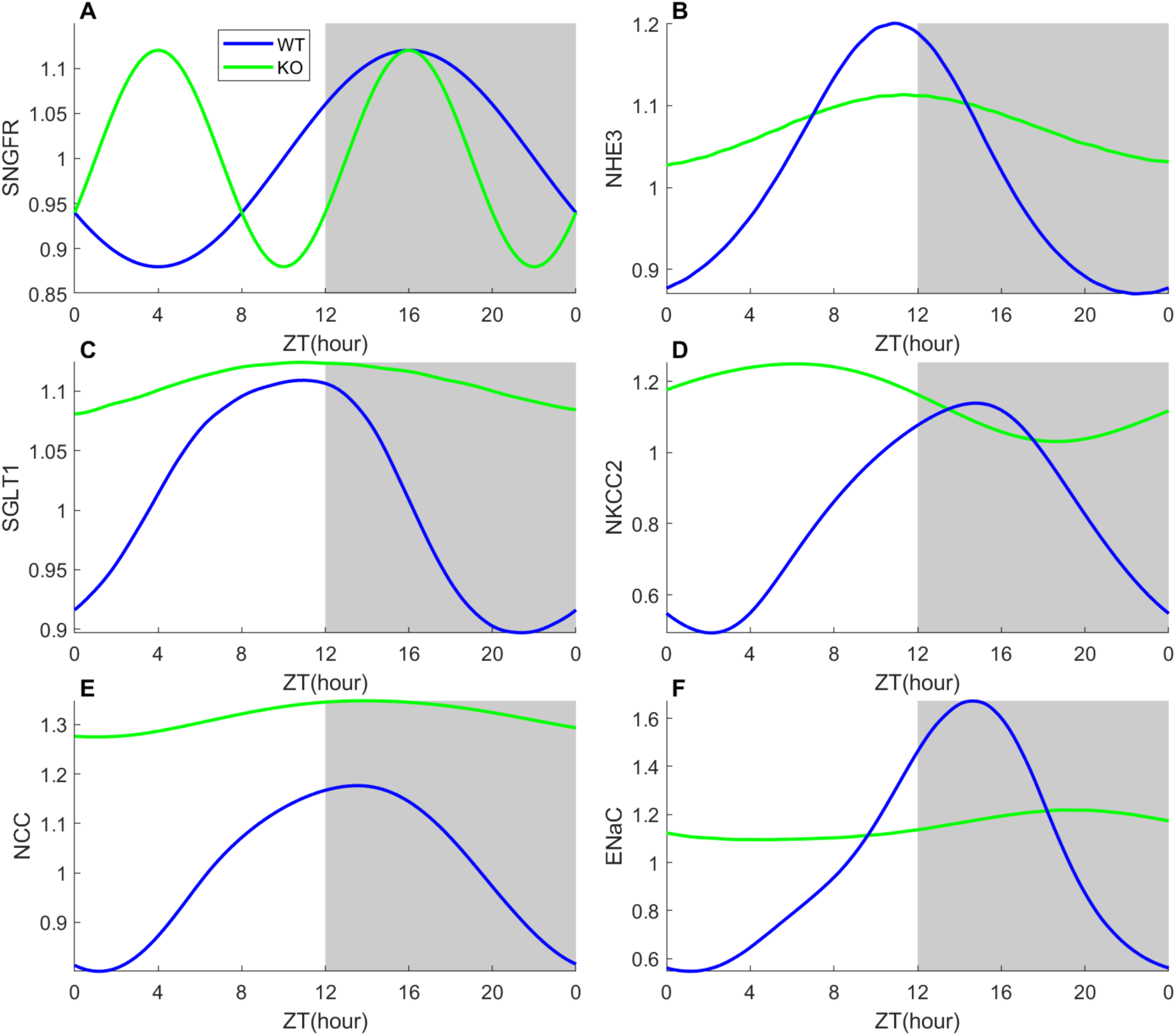
Comparison of model GFR and simulated transporter expression between wild type (WT) and conditional Bmal1 knockout (KO) male rat. (A): GFR is normalized by baseline GFR at ZT10 in the wild type; (B-F) simulated transporter expression levels are normalized by mean experimental values the same as in Fig. 3. Gray shading and white regions correspond to the dark and light phases, respectively.

**Figure 5.**
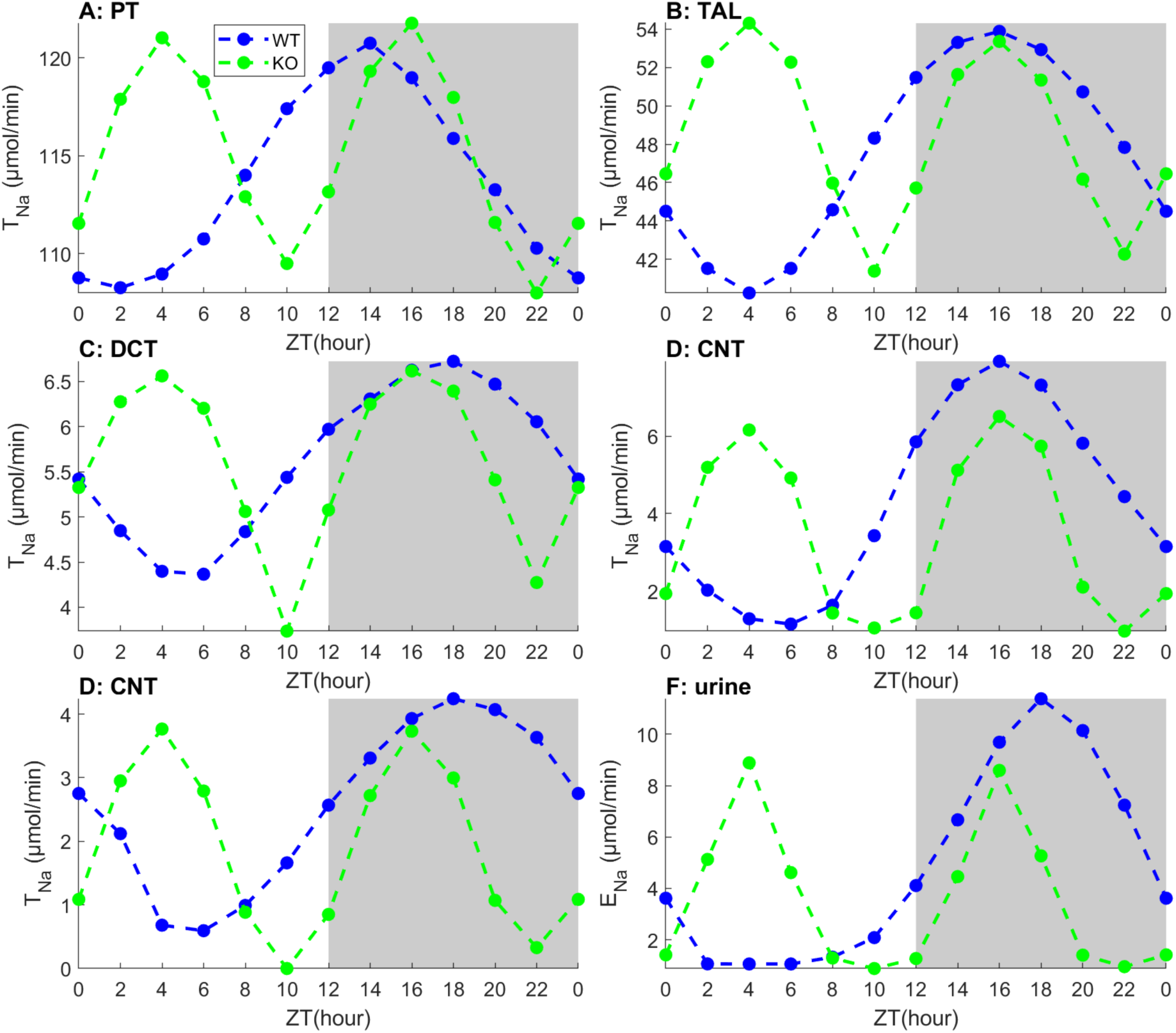
Comparison of segmental Na⁺ transport and urinary Na^+^ in wild type (WT) and Bmal1 knockout (KO) male rats. PT, proximal tubule; TAL, thick ascending limb; DCT, distal convoluted tubule; CNT, connecting tubule; CD, collecting duct. T_Na_, Na⁺ transport; E_Na_, excretion.

Bmal1 knockout leads to a significant reduction in circadian oscillation amplitudes in CLOCK-BMAL1 (results not shown), in NKCC2 as previously noted, and in all other transporters (Fig. 4). The mean transporter expression levels over 24 hours significantly increase, except for NHE3 and ENaC, consistent with the assumption that CLOCK-BMAL1 inhibits these transporters (Fig. 4). Mean NHE3 and ENaC expression levels remain close to wild type. This result may be attributed to the non-linearity of the model, where the mean expressions of NHE3 and ENaC are initially less sensitive to Bmal1 knockout due to the high CLOCK-BMAL1 regulation threshold (Appendix, Table A4). When the maximal Bmal1 transcription rate is reduced to zero, oscillations for all transporters disappear, with the mean expressions of NHE3 and ENaC reduced by 24% and 65%, respectively, relative to wild type.

A comparison between SNGFR in the model wild-type and Bmal1-knockout rats (Fig. 4A) and the predicted urinary Na^+^ excretion (Fig. 5F) suggests that Na^+^ excretion is primarily driven by SNGFR. In Bmal1-knockout rats, cumulative Na^+^ excretion during the inactive (light) phase is 110% higher than wild-type. This difference is primarily due to the higher cumulative SNGFR and thus filtered Na^+^ load in Bmal1-knockout rats in the inactive phase (by 7%). In contrast, during the active (dark) phase, the cumulative SNGFR is 6% lower in Bmal1-knockout rats, and perhaps more importantly, their cumulative activity of NHE3 is 10% higher. As a result, cumulative Na^+^ excretion in Bmal1-knockout rats is predicted to be 55% lower than wild-type. This pattern aligns with experimental findings (40), in which cumulative Na^+^ excretion in Bmal1-knocout male mice was found to be 46% higher than wild type during inactive phase but 24% lower than in active phase.

### Predicted segmental transport and urinary excretions in normotension and hypertension

In the next set of simulations, we assess the differences in nephron function in a normotensive and hypertensive rat throughout the day. Simulations were conducted for a normotensive and a hypertensive rat kidney, using the same SNGFR and clock gene profiles. The predicted filtered Na^+^ load, urinary output, Na^+^ and K^+^ excretions are shown in Fig. 6. Selected predicted segmental transport profiles are shown in Fig. 7.

**Figure 6.**
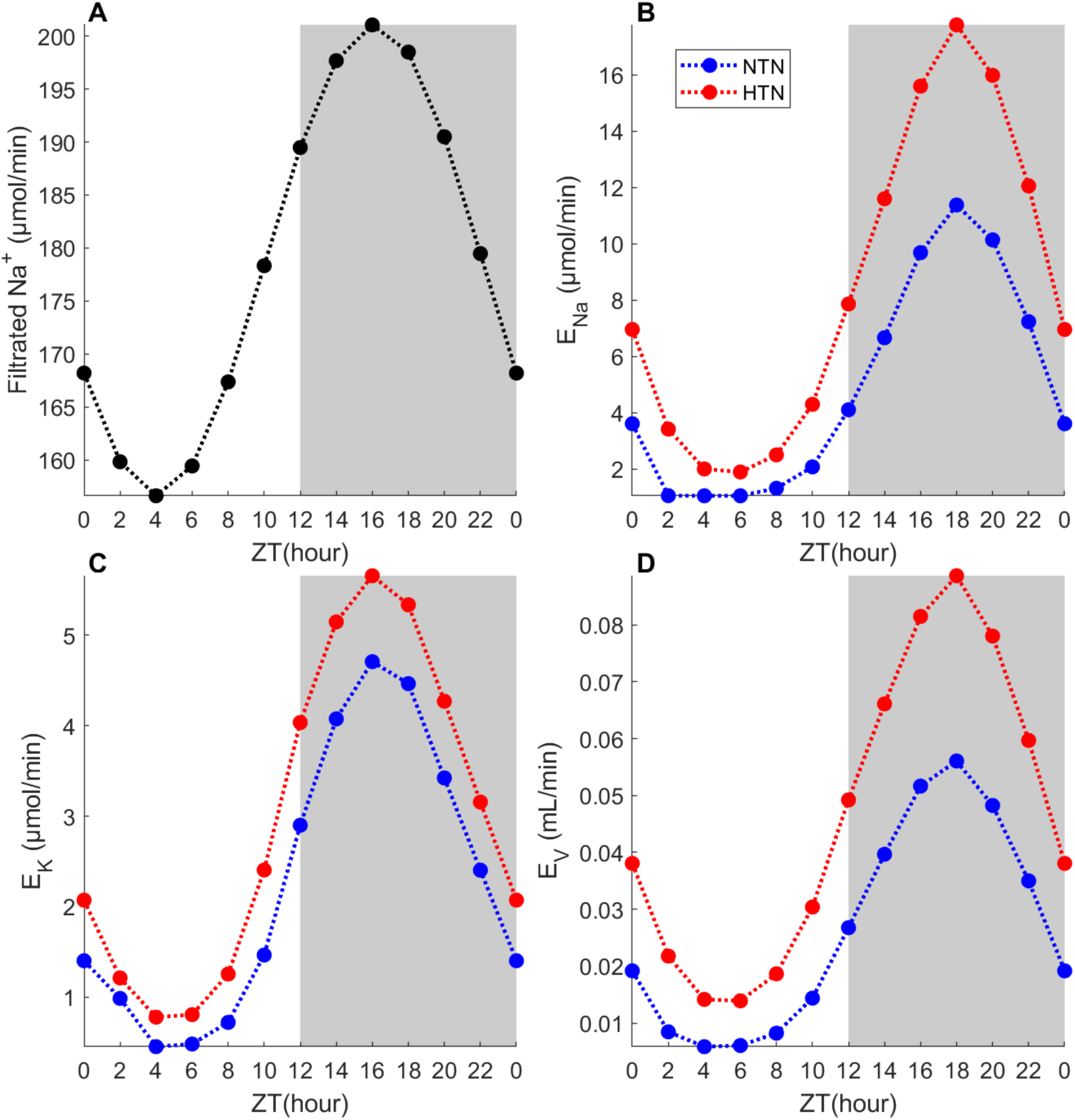
Na⁺ inflow and Na⁺, K⁺, and water excretions. ***A***, Estimated Na⁺ inflow into the nephron for our model under both normotension and hypertension. ***B***-***D***, Na⁺, K⁺, and water excretions under normotension (NTN) and hypertension (HTN). E_Na_, Na⁺ excretion; E_K_, K⁺ excretion; E_V_, water excretion.

**Figure 7.**
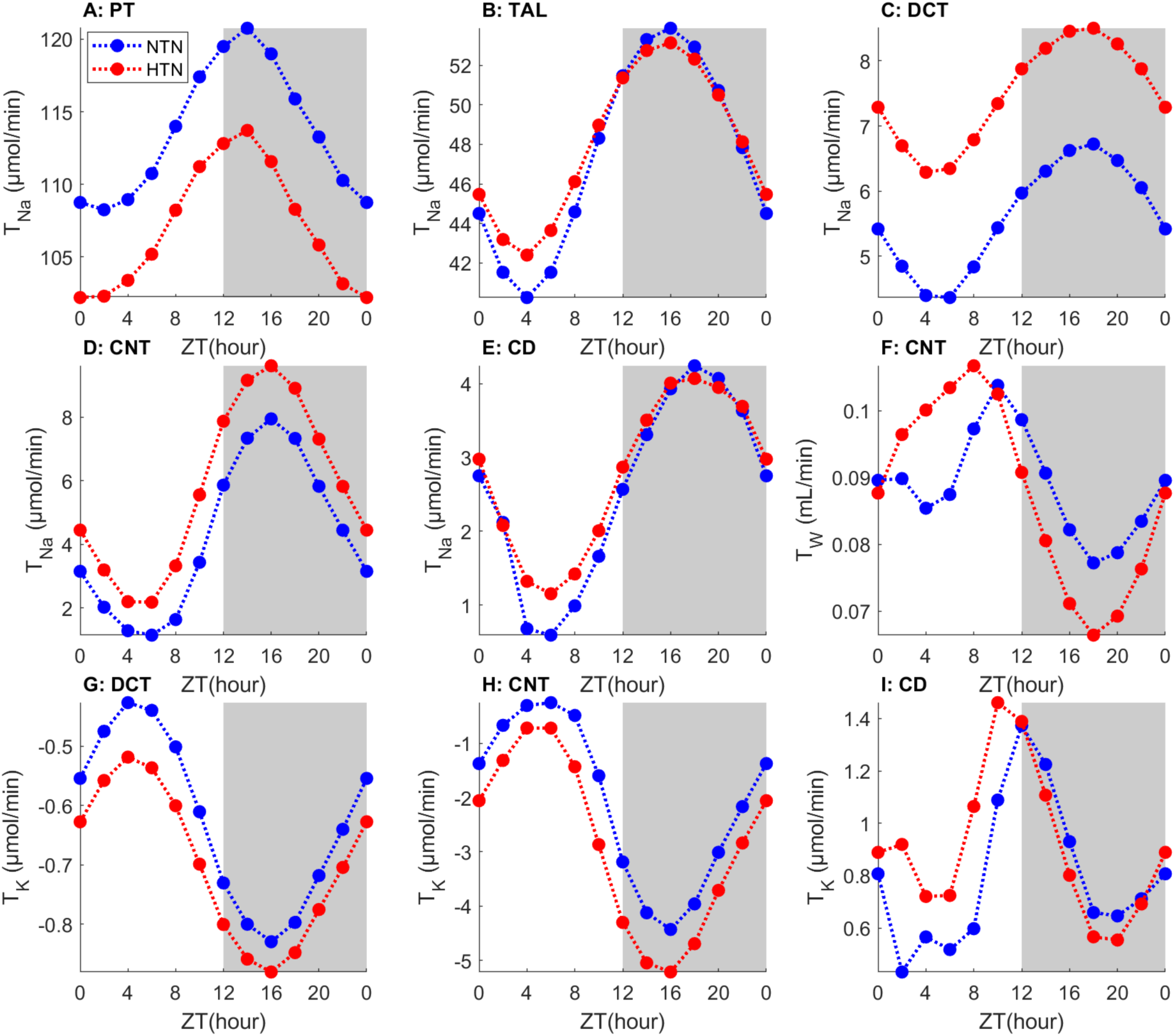
Comparison of segmental Na⁺, K⁺, and water transport under normotension (NTN) and hypertension (HTN). PT, proximal tubule; TAL, thick ascending limb; DCT, distal convoluted tubule; CNT, connecting tubule; CD, collecting duct. T_Na_, Na⁺ transport; T_K_, K⁺ transport; T_W_, water transport.

As expected, the hypertensive-induced downregulation of NHE3 results in reduced Na^+^ reabsorption along the proximal tubule (Fig. 7A). In both normotension and hypertension, proximal tubule Na^+^ reabsorption peaks around ZT14, a result of the competing effects of the diurnal rhythms in SNGFR and NHE3, which peak at ZT16 and ZT11, respectively. Proximal tubule outflow and thus Na^+^ delivery to the loop of Henle downstream peak at ZT16; as a result, Na^+^ reabsorption along the thick ascending limb also peaks at ZT16 (Fig. 7B). Na^+^ transport along the thick ascending limb is similar in normotension and hypertension throughout the day, because the hypertension-induced downregulation of NHE3, NKCC2, and Na^+^-K^+^-ATPase along the medullary thick ascending limb, and the resulting decrease in Na^+^ reabsorption, is compensated by the upregulation of NKCC2 along the cortical thick ascending limb.

Along the downstream segments, Na^+^ reabsorption and K^+^ transport (secretion or reabsorption, depending on the segment) are enhanced by the upregulation of NCC and ENaC in hypertension (Figs. 7C-7I), with the exception of the collecting duct, a result that is consistent with published modeling results (38). That increase persists throughout the day. It is noteworthy that model simulations did not predict a significant phase shift in the segmental transport profiles in hypertension. This result is attributable, in large part, to the assumption of the same diurnal oscillations in SNGFR in normotension and hypertension, which yield similarly shaped oscillations in Na^+^ transport in all segments and in urinary Na^+^ excretion.

Changes in SNGFR and transporter activities together yield oscillations in urine output and excretion rates that are essentially in phase with SNGFR rhythms; see Fig. 6. Urine output and excretions are elevated in hypertension, with larger increases during the dark (active) phase. At ZT4, urine output, Na^+^, K^+^, and water excretions are 90%, 72%, and 140%, respectively, higher in hypertension; at ZT16, the increases are 61%, 20%, and 58%, respectively. It is noteworthy that urinary Na^+^ and volume peak at ZT18, a consequence of high SNGFR and low Na^+^ transporter activities. The predicted patterns of Na⁺ and water excretions align with experimental reports for normotensive and hypertensive mice (41).

### Diuretics induce larger natriuretic and diuretic e7ects during the active phase

Model simulations are then conducted to investigate the effects of administration time of loop, thiazide, and K^+^-sparing diuretics on the function of a hypertensive rat kidney. Predicted segmental Na^+^ transport and urinary excretion are shown in Fig. 8 for ZT4 and ZT16, which correspond to the trough and peak times in the simulated SNGFR. Analogous results for K^+^ and water are shown in Figs. 9 and 10, respectively. As expected, each class of diuretics exerts the largest effect on Na^+^ reabsorption along the primary segment that expresses the target transport: thick ascending limb for loop diuretics, distal convoluted tubule for thiazide diuretics, and connecting tubule and collecting duct for K^+^-sparing diuretics. The lowered Na^+^ reabsorption enhances natriuresis throughout the day, with the effect being larger during the active (dark) phase for all three diuretics. The dark phase is also when Na^+^ reabsorption is higher in each of these segments; see Figs. 8B-8E. The ratio between urinary Na^+^ excretion at ZT16 and ZT4 is approximately 6 for three diuretics (Fig. 8). The natriuretic effect is the largest for the K^+^-sparing diuretics, for both ZT4 and ZT16 (Fig. 8F) because there are fewer segments downstream of its target site (connecting tubule and collecting duct) to compensate for the suppressed ENaC-mediated Na^+^ reabsorption.

**Figure 8.**
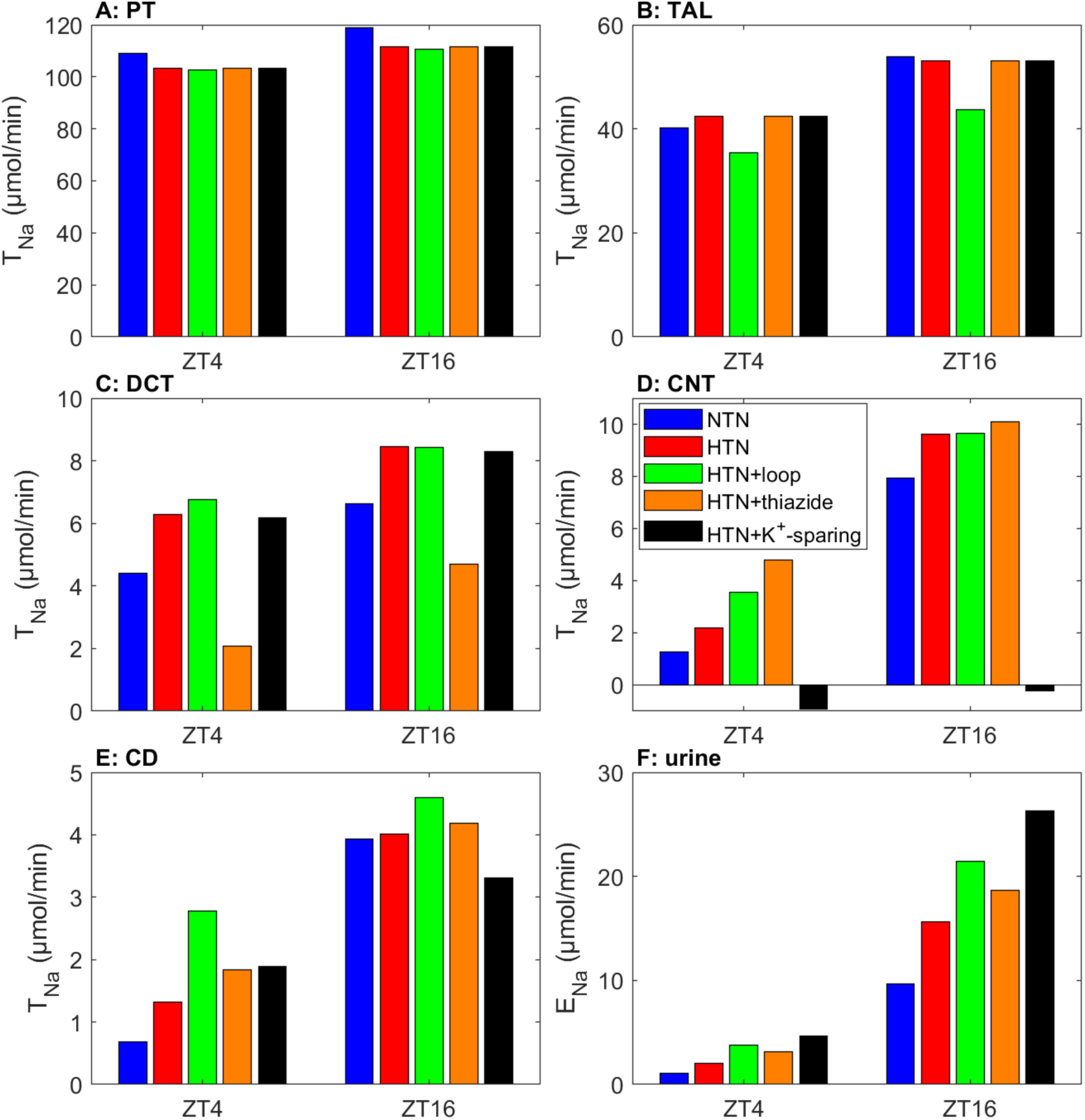
Comparison of Na⁺ transport and excretion, obtained for ZT4 and ZT16, in normotension (NTN), hypertension (HTN), hypertension with loop diuretic administration (HTN+loop), hypertension with thiazide administration (HTN+thiazide), and hypertension with K⁺-spring administration (HTN+K⁺-sparing). PT, proximal tubule; TAL, thick ascending limb; DCT, distal convoluted tubule; CNT, connecting tubule; CD, collecting duct. T_Na_, Na⁺ transport; E_Na_, Na⁺ excretion.

**Figure 9.**
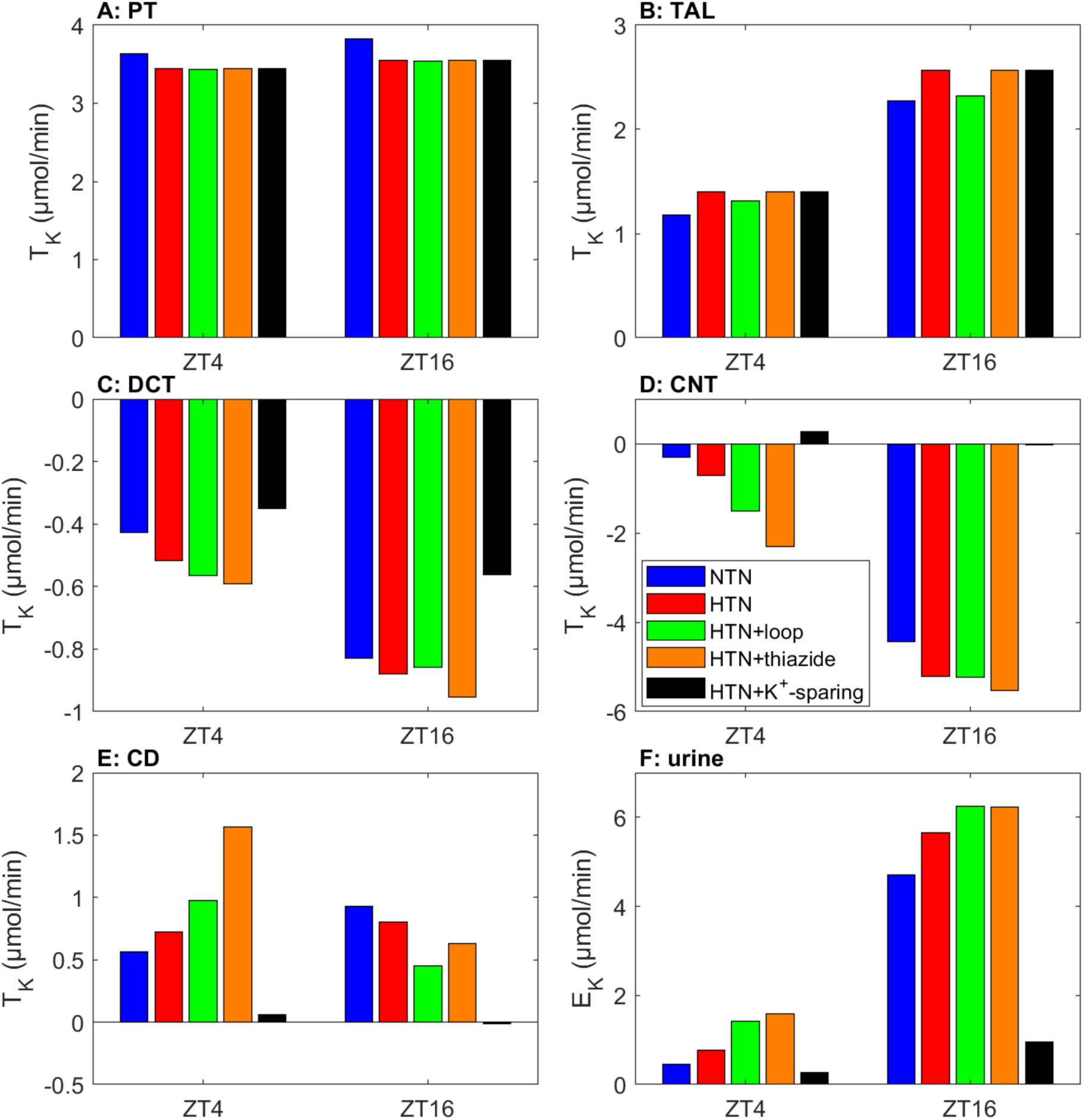
Comparison of K⁺ transport and excretion, obtained for ZT4 and ZT16, in normotension (NTN), hypertension (HTN), hypertension with loop diuretic administration (HTN+loop), hypertension with thiazide administration (HTN+thiazide), and hypertension with K⁺-spring administration (HTN+K⁺-sparing). PT, proximal tubule; TAL, thick ascending limb; DCT, distal convoluted tubule; CNT, connecting tubule; CD, collecting duct. T_K_, K⁺ transport; E_K_, K⁺ excretion.

**Figure 10.**
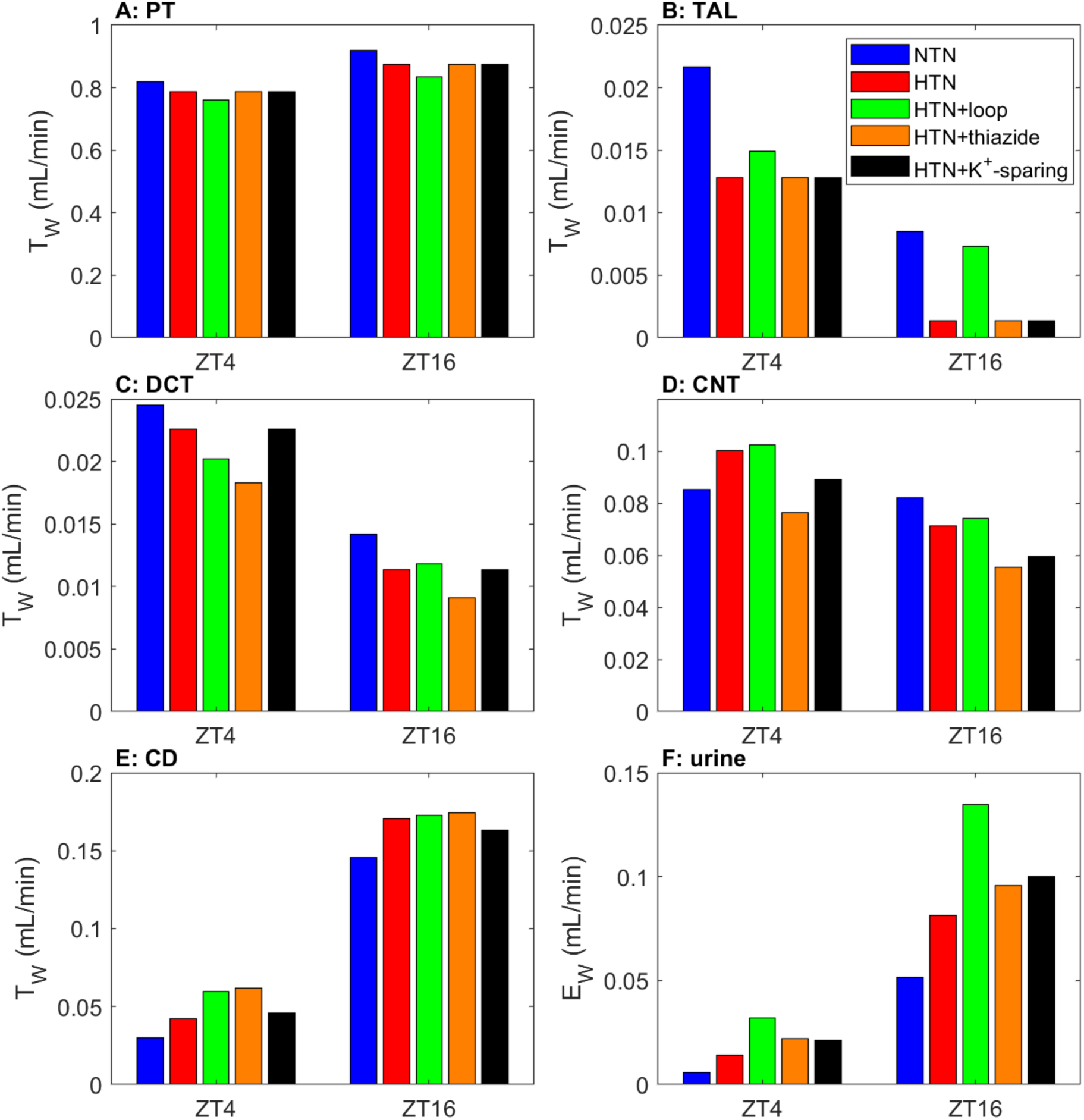
Comparison of water transport and excretion, obtained for ZT4 and ZT16, in normotension (NTN), hypertension (HTN), hypertension with loop diuretic administration (HTN+loop), hypertension with thiazide administration (HTN+thiazide), and hypertension with K⁺-spring administration (HTN+K⁺-sparing). PT, proximal tubule; TAL, thick ascending limb; DCT, distal convoluted tubule; CNT, connecting tubule; CD, collecting duct. T_W_, water transport; E_W_, water excretion.

Consistent with Na^+^ excretion, predicted urinary K^+^ excretion is higher during the dark (active) phase, in both normotension and hypertension, with and without diuretics. Loop and thiazide diuretics both enhance K^+^ excretion, with the increases (but not the actual excretions) somewhat larger during the light (inactive) phase. K^+^-sparing diuretics essentially eliminate K^+^ secretion along the CNT, in fact switching it to a small amount of net reabsorption during the light phase. As intended, K^+^-sparing diuretics lowers K^+^ excretion throughout the day, with the effect larger at ZT16 (-83%) than at ZT4 (-65%); see Fig. 9F.

Water transport essentially follows Na^+^ transport along water-permeable segments. Thus, the predicted segmental water transport and urine output results are qualitatively similar to Na^+^, with the diuretic effect predicted to be larger during the active (dark) phase; see Fig. 10. The diuretic effect is largest for loop diuretics, because it impairs the concentrating mechanism and lowers interstitial fluid osmolality, resulting in attenuated water reabsorption and elevated urine volume (Fig. 10F). This is true for both normotension and hypertension.

## DISCUSSION

Even though the occurrence of circadian rhythms is well known in several renal functions (42,43), their clinical importance has yet to be completely understood. In both humans and rodents, disruption of the circadian rhythms, including changes in time zone, is associated with an increased risk of cardiovascular disease (44,45). Given the marked diurnal variations in urinary excretion, the plasma concentration or bioavailability of drugs that are eliminated by the kidneys might depend on the circadian expression of renal transporters, even if these drugs do not target renal transporters. The circadian rhythm is also known to affect the pathogenesis or progression of some renal diseases, such as renal tissue fibrosis (46) and kidney stones (47,48). The morbidity of patients with chronic kidney disease, as well as patients on hemodialysis or peritoneal dialysis, may be partially attributed to the disruption of the circadian rhythm of key renal functions (49,50). The circadian rhythm is followed by blood pressure in humans as well, with values 10–15% lower during nighttime than during daytime, in a phenomenon known as dipping (51). The absence of a nocturnal blood pressure decrease (i.e., non-dipping) is associated with target organ damage (52). The interactions among the central and peripheral clocks, and the organ and tissue systems, are intriguing and complex. What are the underlying mechanisms that lead to the emergence of circadian rhythms in these processes? For example, the determinants of blood pressure dipping and its disappearance has yet to be understood. What are the clinical implications of the disappearance of these circadian rhythms? And how should pharmacological treatments be timed to maximize their effectiveness?

As a step towards answering these questions, we developed a detailed computational model of the circadian regulation of epithelial transport in a rat kidney. Most published modeling studies of nephron epithelial transport focus on steady-state results (6,38,39,53,54), with a few exceptions such as Refs. (36,55–57), where the circadian regulation of key renal transporters is represented. A key difference between those published studies (36,55) and the present model is that the former represent mouse kidneys, whereas the present model simulates a rat kidney. Also, the published studies (36,55–57) prescribe the circadian oscillations of the transporter activities, whereas in the present study, transporter activities are computed based on circadian protein levels, which are predicted by a detailed computational model of the renal circadian clock that simulates the interactions among Per, Cry, Bmal1, Clock, Ror, and Rev-Erb. By modeling the clock explicitly, we are able to predict the effect of clock gene knockout on kidney function (see Fig. 1).

By modeling the renal circadian clock and its regulation of transporter activities, our model can predict the extent to which Na^+^, K^+^ and water transport by a nephron is modulated by the renal circadian clock. A similar renal circadian clock was developed in Ref. (58) to predict the circadian regulation of Na^+^ and water transport by a proximal convoluted tubule cell. That model (58) does not differentiate between homologs of Per and Cry. However, Per1 and Per2 have been shown to exert opposite effects on NHE3 expression (25,26). Even though direct evidence is not currently available, it does not seem unreasonable to imagine that distinct roles of Per and Cry homologs on the circadian regulation of key renal transporters may be revealed in the future. Thus, the present model represents the Per and Cry homologs separately and allows the four protein complexes (PER1-CRY1, PER1-CRY2, PER2-CRY1, PER2-CRY2) to have different effects on transporter expression levels.

Emans et al. (59) reported that wild-type rats exhibit a circadian rhythm in kidney tissue oxygenation, with lower levels during the light phase and higher levels during the dark phase, peaking at ZT16.9. Kidney oxygenation is a result of the balance of oxygen delivery and consumption, which in turn are determined by renal hemodynamics and metabolism, respectively. In contrast to the 24-hour circadian rhythm in wild-types, SNGFR and thus Na^+^ transport in Bmal1-knockout exhibit a double-peak, 12-hour rhythm (Fig. 4A). Given that renal metabolism is driven primarily by tubular active Na^+^ transport, we hypothesize that renal tissue oxygenation in Bmal1-knockout exhibits a different rhythm, with a 12-hour period instead of 24.

In Ref. (38), we published the first computational model of epithelial transport in the kidney of a rat with hypertension. That model was formulated for steady state. With circadian rhythms incorporated, the present model predicts that due to pressure natriuresis, Na^+^, K^+^, and water transport is suppressed along the proximal tubule in the hypertensive rat, but is generally elevated along the distal segments (Fig. 7). These competing effects result in hypertension-enhanced natriuresis, diuresis, and kaliuresis throughout the day (Fig. 6). The hypertensive kidney model allows us to investigate how administration time may impact the natriuretic and diuretic effects of diuretics in a rat with hypertension. In fact, while Dutta et al. simulated the administration of the same three classes of diuretics (55), they used a normotensive kidney. Our simulations indicate that effects of the diuretics are qualitatively similar in the normotensive and hypertensive kidneys (Fig. 8-10). All three diuretics yield stronger natriuretic and diuretic effects during the active phase, with K^+^-sparing and loop diuretics producing the strongest natriuretic and diuretic effects, respectively (Figs. 8 and 9).

The effectiveness of an antihypertensive medication is typically measured by how much it lowers blood pressure. The kidney model used in the present study predicts urine output and excretions, but not extracellular fluid volume or blood pressure. Indeed, the present model does not include other key regulatory mechanisms in blood pressure regulation, such as the renin-angiotensin system, renal sympathetic nervous system, oxidative stress/nitric oxide pathway, and endothelin system. Some of these systems regulate blood pressure by acting on the kidneys. Furthermore, circadian oscillations in GFR and urinary excretion may impact the drug’s excretion and pharmacokinetics. Thus, while diuresis and natriuresis, taken in isolation, would lower blood pressure, the picture is complicated by the contributions of the other systems noted above. To assess the time-of-day variations of the effectiveness of antihypertensive drugs, the present nephron models can be combined with whole-body blood pressure regulation models (60,61). The resulting integrative model would take into account the multiple feedback controls and correctly link natriuresis and diuresis to extracellular fluid volume and blood pressure.

Besides the circadian clock, sex is another key modulator of blood pressure regulation. Sex-based differences in blood pressure and its dysregulation have been observed across various mammalian and avian species (62). Studies in animal models of hypertension, such as spontaneously hypertensive rats (63) and Dahl salt-sensitive rats (64), reveal that males tend to develop hypertension earlier and more severely than females. Similarly, premenopausal women typically exhibit lower blood pressure and a lower incidence of hypertension compared to men of the same age (65), although that female advantage vanishes in postmenopausal women over the age of 60. Sex differences are evident across organ and tissue systems that are involved in blood pressure regulation, collectively contributing to differences in blood pressure and hypertension between the sexes. The present model simulates kidney function in a male rat. Future studies should extend the present model to a female rat kidney, followed by whole-body models of circadian regulation of blood pressure control in both sexes. Model simulations may yield insights into the mechanisms via which each transcription factor contributes to blood pressure phenotype, and explain any sex differences.

## Appendix: Equations and Parameters

### A1 Governing Equations Overview

Eq. A1-A21 are 25 governing equations of 12 mRNAs: Per1, Per2, Cry1, Cry2, Rev-Erb, Ror, Bmal1, NHE3, NKCC2, SGLT1,NCC, and ENaC; 7 proteins: PER1, PER2, CRY1, CRY2, REV-ERB, ROR, and BMAL1; and 5 protein complexes: PER1-CRY1, PER2-CRY1, PER1-CRY2, PER2-CRY2, and CLOCK-BMAL1. Because in our formulation all mRNA are regualted by 5 protein complexes in an analogous pattern, we unify the formulation by defining a regulation factor F_X_ (Eq. A13) where X is the name of mRNA. All parameters are listed with description in Section A4.

### A2 mRNA equations

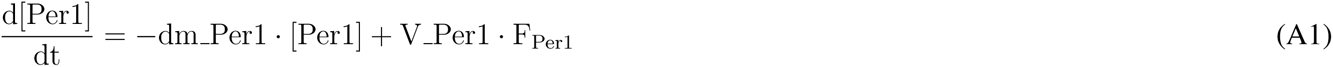

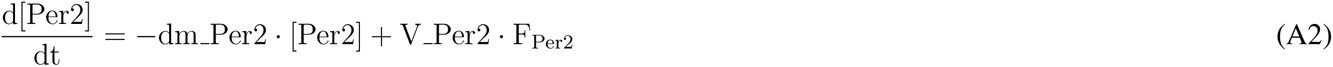

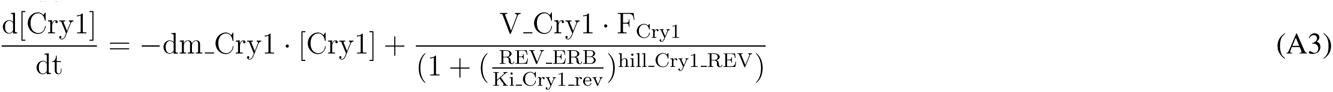

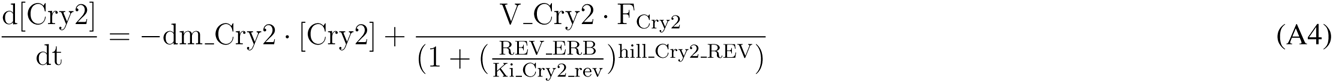

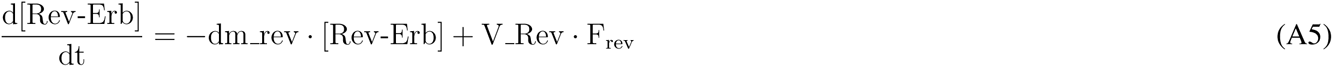

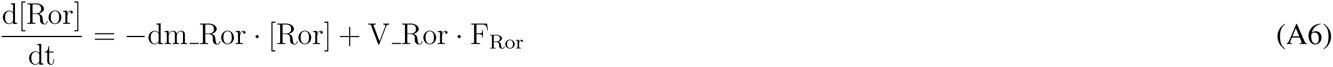

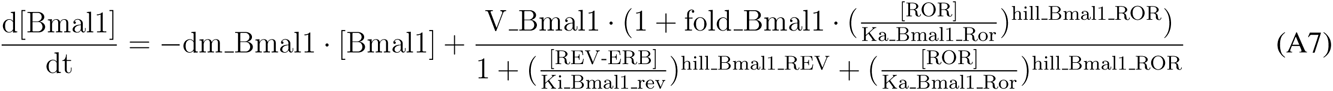

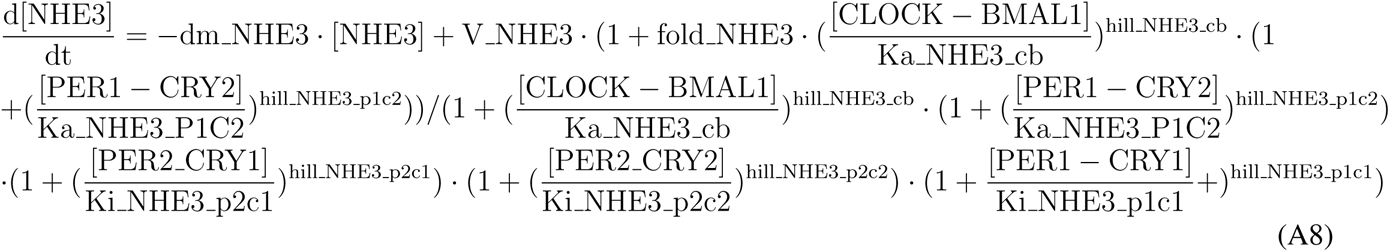

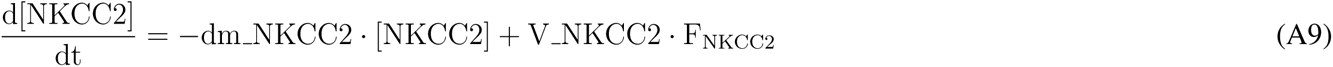

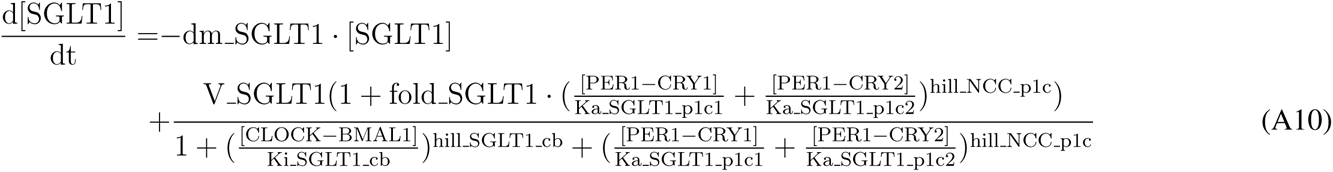

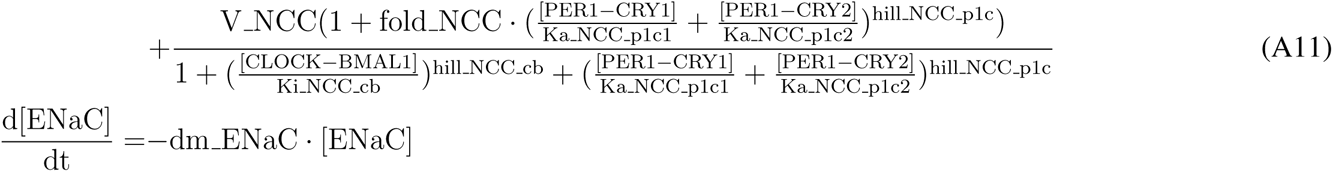

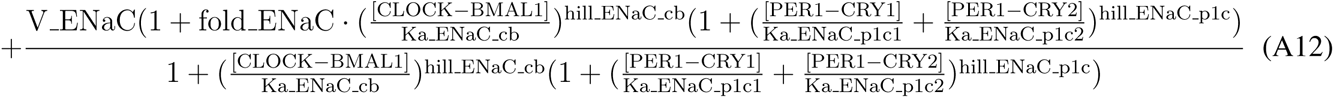

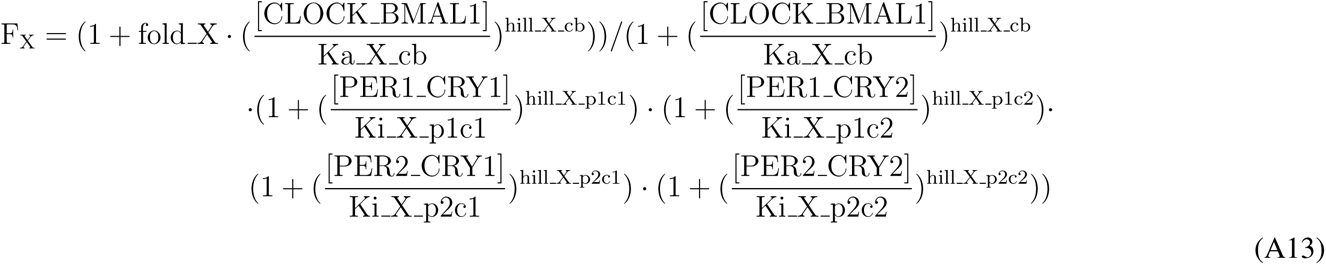

### A3 Protein Equations

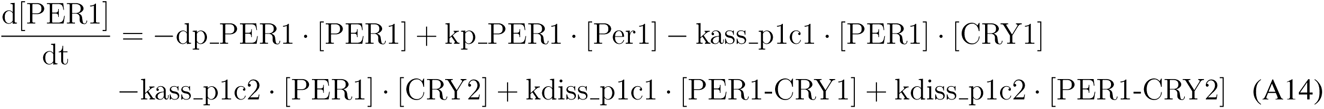

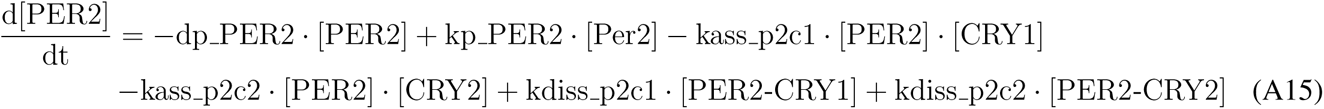

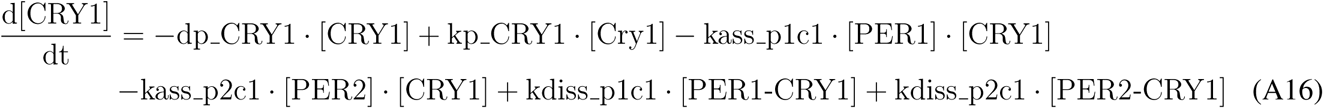

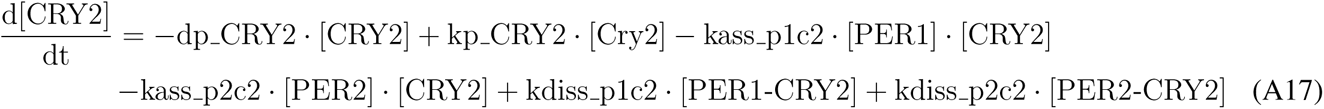

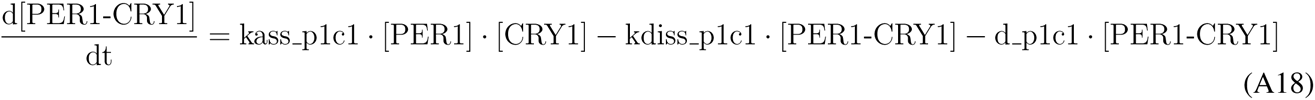

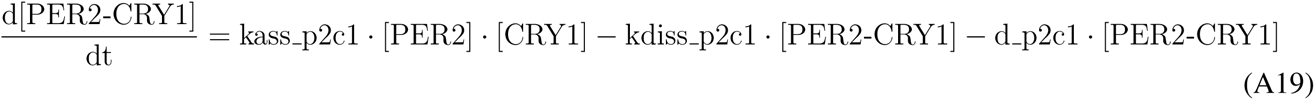

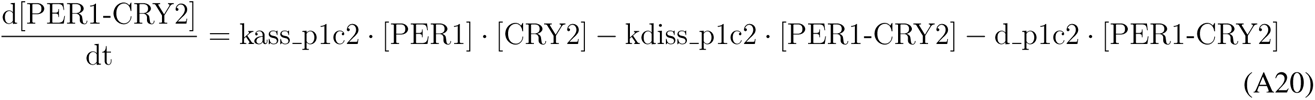

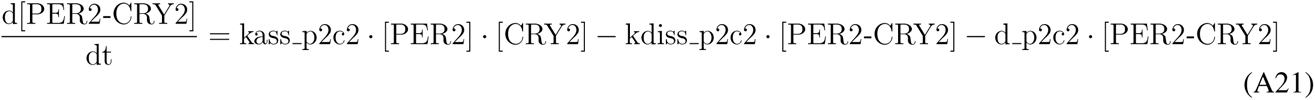

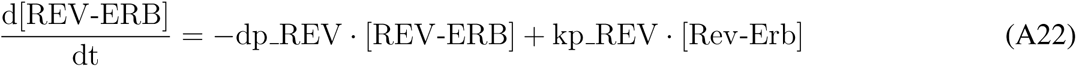

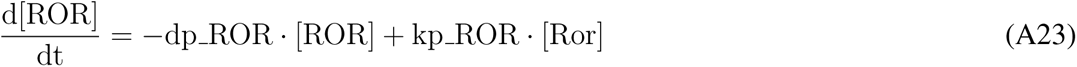

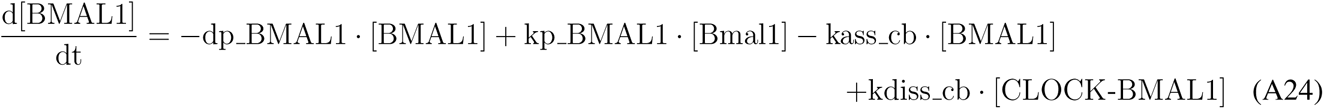

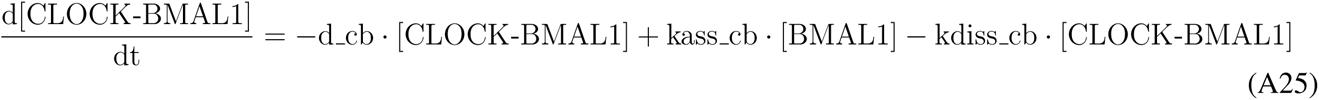

### A4 Baseline parameters

**Table A1:** Degradation rates (in hour*^−^*^1^) of protein complexes, mRNAs, and proteins.

| Parameter | Value | Description |
| --- | --- | --- |
| d_cb | 0.098775151 | CLOCK-BMAL1 complex degradation rate |
| d_p1c1 | 2.747914868 | PER1-CRY1 protein complex degradation rate |
| d_p1c2 | 3.857884688 | PER1-CRY2 protein complex degradation rate |
| d_p2c1 | 1.98150769 | PER2-CRY1 protein complex degradation rate |
| d_p2c2 | 0.009962143 | PER2-CRY2 protein complex degradation rate |
| dm_Bmal1 | 10.29999143 | Bmal1 mRNA degradation rate |
| dm_Cry1 | 1.680357077 | Cry1 mRNA degradation rate |
| dm_Cry2 | 1.102297307 | Cry2 mRNA degradation rate |
| dm_Per1 | 0.393284589 | Per1 mRNA degradation rate |
| dm_Per2 | 0.199538876 | Per2 mRNA degradation rate |
| dm_Rev | 0.202152455 | Rev mRNA degradation rate |
| dm_Ror | 0.733726737 | Ror mRNA degradation rate |
| dm_NHE3 | 2.6212974828 | NHE3 mRNA degradation rate |
| dm_NKCC2 | 0.147706095 | NKCC2 mRNA degradation rate |
| dm_SGLT1 | 0.9719928901 | SGLT1 mRNA degradation rate |
| dm_NCC | 0.11428499820 | NCC mRNA degradation rate |
| dm_ENaC | 1.4666546516 | ENaC mRNA degradation rate |
| dp_BMAL1 | 0.1223564 | BMAL1 protein degradation rate |
| dp_CRY1 | 0.07170199 | CRY1 protein degradation rate |
| dp_CRY2 | 0.063886143 | CRY2 protein degradation rate |
| dp_PER1 | 2.118148503 | PER1 protein degradation rate |
| dp_PER2 | 1.092137167 | PER2 protein degradation rate |
| dp_REV | 1.54249893 | REV-ERB protein degradation rate |
| dp_ROR | 0.110251246 | ROR protein degradation rate |

**Table A2:** Activation ratios (dimensionless).

| Parameter | Value | Description |
| --- | --- | --- |
| fold_Bmal1 | 29.23079974 | Activation ratio of Bmal1 by ROR |
| fold_Cry1 | 17.45987491 | Activation ratio of Cry1 by CLOCK-BMAL1 |
| fold_Cry2 | 13.21473162 | Activation ratio of Cry2 by CLOCK-BMAL1 |
| fold_Per1 | 2.431717595 | Activation ratio of Per1 by CLOCK-BMAL1 |
| fold_Per2 | 3.308354134 | Activation ratio of Per2 by CLOCK-BMAL1 |
| fold_Rev | 118.5947203 | Activation ratio of Rev by CLOCK-BMAL1 |
| fold_Ror | 5.603448203 | Activation ratio of Ror by CLOCK-BMAL1 |
| fold_NHE3 | 5.63209402975596 | Activation ratio of NHE3 by CLOCK-BMAL1 and PER1-CRY2 |
| fold_NKCC2 | 0.385311524 | Activation ratio of NKCC2 by CLOCK-BMAL1 |
| fold_SGLT1 | 2.36457412221910 | Activation ratio of SGLT1 by CLOCK-BMAL1 |
| fold_NCC | 1.026401864136 | Activation ratio of NCC by PER1-CRY1 and PER1-CRY2 |
| fold_ENaC | 6.2735611435 | Activation ratio of ENaC by CLOCK-BMAL1 |

**Table A3:** Hill coefficients (dimensionless).

| Parameter | Value | Description |
| --- | --- | --- |
| hill_Bmal1_REV | 6.315290784 | Hill coefficient regulation of Bmal1 by REV-ERB |
| hill_Bmal1_ROR | 10.19219221 | Hill coefficient regulation of Bmal1 by ROR |
| hill_Cry1_cb | 4.565210968 | Hill coefficient regulation of Cry1 by CLOCK-BMAL1 |
| hill_Cry1_p1c1 | 1.768766266 | Hill coefficient regulation of Cry1 by PER1-CRY1 |
| hill_Cry1_p1c2 | 1.438000996 | Hill coefficient regulation of Cry1 by PER1-CRY2 |
| hill_Cry1_p2c1 | 1.589908884 | Hill coefficient regulation of Cry1 by PER2-CRY1 |
| hill_Cry1_p2c2 | 1.830362004 | Hill coefficient regulation of Cry1 by PER2-CRY2 |
| hill_Cry1_Rev | 4.820086463 | Hill coefficient regulation of Cry1 by REV-ERB |
| hill_Cry2_cb | 1.367823416 | Hill coefficient regulation of Cry2 by CLOCK-BMAL1 |
| hill_Cry2_p1c1 | 2.1098479 | Hill coefficient regulation of Cry2 by PER1-CRY1 |
| hill_Cry2_p1c2 | 2.026033401 | Hill coefficient regulation of Cry2 by PER1-CRY2 |
| hill_Cry2_p2c1 | 1.602187775 | Hill coefficient regulation of Cry2 by PER2-CRY1 |
| hill_Cry2_p2c2 | 1.655287085 | Hill coefficient regulation of Cry2 by PER2-CRY2 |
| hill_Cry2_rev | 1.032736435 | Hill coefficient regulation of Cry2 by REV-ERB |
| hill_Per1_cb | 4.096754782 | Hill coefficient regulation of Per1 by CLOCK-BMAL1 |
| hill_Per1_p1c1 | 4.137208965 | Hill coefficient regulation of Per1 by PER1-CRY1 |
| hill_Per1_p1c2 | 4.352895626 | Hill coefficient regulation of Per1 by PER1-CRY2 |
| hill_Per1_p2c1 | 4.556232531 | Hill coefficient regulation of Per1 by PER2-CRY1 |
| hill_Per1_p2c2 | 3.006414519 | Hill coefficient regulation of Per1 by PER2-CRY2 |
| hill_Per2_cb | 5.973526978 | Hill coefficient regulation of Per2 by CLOCK-BMAL1 |
| hill_Per2_p1c1 | 4.917946201 | Hill coefficient regulation of Per2 by PER1-CRY1 |
| hill_Per2_p1c2 | 4.094774755 | Hill coefficient regulation of Per2 by PER1-CRY2 |
| hill_Per2_p2c1 | 4.287994314 | Hill coefficient regulation of Per2 by PER2-CRY1 |
| hill_Per2_p2c2 | 5.627510726 | Hill coefficient regulation of Per2 by PER2-CRY2 |
| hill_Rev_cb | 13.59046465 | Hill coefficient regulation of Rev-Erb by CLOCK-BMAL1 |
| hill_Rev_p1c1 | 4.593824878 | Hill coefficient regulation of Rev-Erb by PER1-CRY1 |
| hill_Rev_p1c2 | 4.473939935 | Hill coefficient regulation of Rev-Erb by PER1-CRY2 |
| hill_Rev_p2c1 | 4.502871061 | Hill coefficient regulation of Rev-Erb by PER2-CRY1 |
| hill_Rev_p2c2 | 4.474437607 | Hill coefficient regulation of Rev-Erb by PER2-CRY2 |
| hill_Ror_cb | 11.23225359 | Hill coefficient regulation of Ror by CLOCK-BMAL1 |
| hill_Ror_p1c1 | 3.346138809 | Hill coefficient regulation of Ror by PER1-CRY1 |
| hill_Ror_p1c2 | 4.279505041 | Hill coefficient regulation of Ror by PER1-CRY2 |
| hill_Ror_p2c1 | 3.213895403 | Hill coefficient regulation of Ror by PER2-CRY1 |
| hill_Ror_p2c2 | 4.047251298 | Hill coefficient regulation of Ror by PER2-CRY2 |
| hill_NHE3_cb | 2.2098682284 | Hill coefficient regulation of NHE3 by CLOCK-BMAL1 |
| hill_NHE3_p1c1 | 4.0457153929 | Hill coefficient regulation of NHE3 by PER1-CRY1 |
| hill_NHE3_p1c2 | 5.8872697770 | Hill coefficient regulation of NHE3 by PER1-CRY2 |
| hill_NHE3_p2c1 | 5.1885742129 | Hill coefficient regulation of NHE3 by PER2-CRY1 |
| hill_NHE3_p2c2 | 1.7481499102 | Hill coefficient regulation of NHE3 by PER2-CRY2 |
| hill_NKCC2_cb | 21.63335325 | Hill coefficient regulation of NKCC2 by CLOCK-BMAL1 |
| hill_NKCC2_p1c1 | 10.57552098 | Hill coefficient regulation of NKCC2 by PER1-CRY1 |
| hill_NKCC2_p1c2 | 11.3012353 | Hill coefficient regulation of NKCC2 by PER1-CRY2 |
| hill_NKCC2_p2c1 | 11.0258738 | Hill coefficient regulation of NKCC2 by PER2-CRY1 |
| hill_NKCC2_p2c2 | 15.74026414 | Hill coefficient regulation of NKCC2 by PER2-CRY2 |
| hill_SGLT1_cb | 1.8701185743 | Hill coefficient regulation of SGLT1 by CLOCK-BMAL1 |
| hill_SGLT1_p1c | 5.86981584935250 | Hill coefficient regulation of SGLT1 by PER1-CRY1 and PER1-CRY2 |
| hill_NCC_cb | 2.1869392091 | Hill coefficient regulation of NCC by CLOCK-BMAL1 |
| hill_NCC_p1c | 4.8511288196 | Hill coefficient regulation of NCC by PER1-CRY1 and PER1-CRY2 |
| hill_ENaC_cb | 6.4999347889 | Hill coefficient regulation of ENaC by CLOCK-BMAL1 |
| hill_ENaC_p1c | 10.4928793515923 | Hill coefficient regulation of ENaC by PER1-CRY1 and PER1-CRY2 |

**Table A4:** Regulation thresholds (dimensionless).

| Parameter | Value | Description |
| --- | --- | --- |
| Ka_Bmal1_ror | 0.788134198 | Regulation threshold of Bmal1 by ROR |
| Ka_Cry1_cb | 2.965895772 | Regulation threshold of Cry1 by CLOCK-BMAL1 |
| Ka_Cry2_cb | 9.818058078 | Regulation threshold of Cry2 by CLOCK-BMAL1 |
| Ka_Per1_cb | 3.045974034 | Regulation threshold of Per1 by CLOCK-BMAL1 |
| Ka_Per2_cb | 0.996030215 | Regulation threshold of Per2 by CLOCK-BMAL1 |
| Ka_Rev_cb | 2.407381652 | Regulation threshold of Rev-Erb by CLOCK-BMAL1 |
| Ka_Ror_cb | 2.531369156 | Regulation threshold of Ror by CLOCK-BMAL1 |
| Ka_NHE3_cb | 3.0427238397 | Regulation threshold of NHE3 by CLOCK-BMAL1 |
| Ka_NKCC2_cb | 1.238599283 | Regulation threshold of NKCC2 by CLOCK-BMAL1 |
| Ki_SGLT1_cb | 5.414004356 | Regulation threshold of SGLT1 by CLOCK-BMAL1 |
| Ki_NCC_cb | 0.5108645898 | Regulation threshold of NCC by CLOCK-BMAL1 |
| Ka_ENaC_cb | 3.2433974568 | Regulation threshold of ENaC by CLOCK-BMAL1 |
| Ki_Bmal1_rev | 0.000960248 | Regulation threshold of Bmal1 by REV-ERB |
| Ki_Cry1_p1c1 | 2.559548048 | Regulation threshold of Cry1 by PER1-CRY1 |
| Ki_Cry1_p1c2 | 2.536630442 | Regulation threshold of Cry1 by PER1-CRY2 |
| Ki_Cry1_p2c1 | 2.455448644 | Regulation threshold of Cry1 by PER2-CRY1 |
| Ki_Cry1_p2c2 | 3.054560489 | Regulation threshold of Cry1 by PER2-CRY2 |
| Ki_Cry1_rev | 3.597110237 | Regulation threshold of Cry1 by REV-ERB |
| Ki_Cry2_p1c1 | 2.852145929 | Regulation threshold of Cry2 by PER1-CRY1 |
| Ki_Cry2_p1c2 | 3.094816119 | Regulation threshold of Cry2 by PER1-CRY2 |
| Ki_Cry2_p2c1 | 2.709023574 | Regulation threshold of Cry2 by PER2-CRY1 |
| Ki_Cry2_p2c2 | 2.309457939 | Regulation threshold of Cry2 by PER2-CRY2 |
| Ki_Cry2_rev | 0.378297016 | Regulation threshold of Cry2 by REV-ERB |
| Ki_Per1_p1c1 | 1.010242998 | Regulation threshold of Per1 by PER1-CRY1 |
| Ki_Per1_p1c2 | 1.721720548 | Regulation threshold of Per1 by PER1-CRY2 |
| Ki_Per1_p2c1 | 1.624880971 | Regulation threshold of Per1 by PER2-CRY1 |
| Ki_Per1_p2c2 | 0.368879768 | Regulation threshold of Per1 by PER2-CRY2 |
| Ki_Per2_p1c1 | 1.063344333 | Regulation threshold of Per2 by PER1-CRY1 |
| Ki_Per2_p1c2 | 0.568083313 | Regulation threshold of Per2 by PER1-CRY2 |
| Ki_Per2_p2c1 | 0.86599114 | Regulation threshold of Per2 by PER2-CRY1 |
| Ki_Per2_p2c2 | 0.467627765 | Regulation threshold of Per2 by PER2-CRY2 |
| Ki_Rev_p1c1 | 84.46016203 | Regulation threshold of Rev-Erb by PER1-CRY1 |
| Ki_Rev_p1c2 | 83.99580669 | Regulation threshold of Rev-Erb by PER1-CRY2 |
| Ki_Rev_p2c1 | 84.00271945 | Regulation threshold of Rev-Erb by PER2-CRY1 |
| Ki_Rev_p2c2 | 83.99582755 | Regulation threshold of Rev-Erb by PER2-CRY2 |
| Ki_Ror_p1c1 | 1.094009455 | Regulation threshold of Ror by PER1-CRY1 |
| Ki_Ror_p1c2 | 2.291802041 | Regulation threshold of Ror by PER1-CRY2 |
| Ki_Ror_p2c1 | 0.17734771 | Regulation threshold of Ror by PER2-CRY1 |
| Ki_Ror_p2c2 | 3.087808611 | Regulation threshold of Ror by PER2-CRY2 |
| Ki_NHE3_p1c1 | 0.5248622142 | Regulation threshold of NHE3 by PER1-CRY1 |
| Ka_NHE3_p1c2 | 0.51148790424 | Regulation threshold of NHE3 by PER1-CRY2 |
| Ki_NHE3_p2c1 | 0.04273092406 | Regulation threshold of NHE3 by PER2-CRY1 |
| Ki_NHE3_p2c2 | 0.50842406780 | Regulation threshold of NHE3 by PER2-CRY2 |
| Ki_NKCC2_p1c1 | 7.111624431 | Regulation threshold of NKCC2 by PER1-CRY1 |
| Ki_NKCC2_p1c2 | 6.374145136 | Regulation threshold of NKCC2 by PER1-CRY2 |
| Ki_NKCC2_p2c1 | 6.328221462 | Regulation threshold of NKCC2 by PER2-CRY1 |
| Ki_NKCC2_p2c2 | 0.944237431 | Regulation threshold of NKCC2 by PER2-CRY2 |
| Ka_SGLT1_p1c1 | 5.579866217 | Regulation threshold of SGLT1 by PER1-CRY1 |
| Ka_SGLT1_p1c2 | 0.3368079674 | Regulation threshold of SGLT1 by PER1-CRY2 |
| Ka_NCC_p1c1 | 0.61559096740 | Regulation threshold of NCC by PER1-CRY1 |
| Ka_NCC_p1c2 | 0.17552314947 | Regulation threshold of NCC by PER1-CRY2 |
| Ka_ENaC_p1c1 | 1.0303132234 | Regulation threshold of ENaC by PER1-CRY1 |
| Ka_ENaC_p1c2 | 0.22547393780 | Regulation threshold of ENaC by PER1-CRY2 |

**Table A5:**
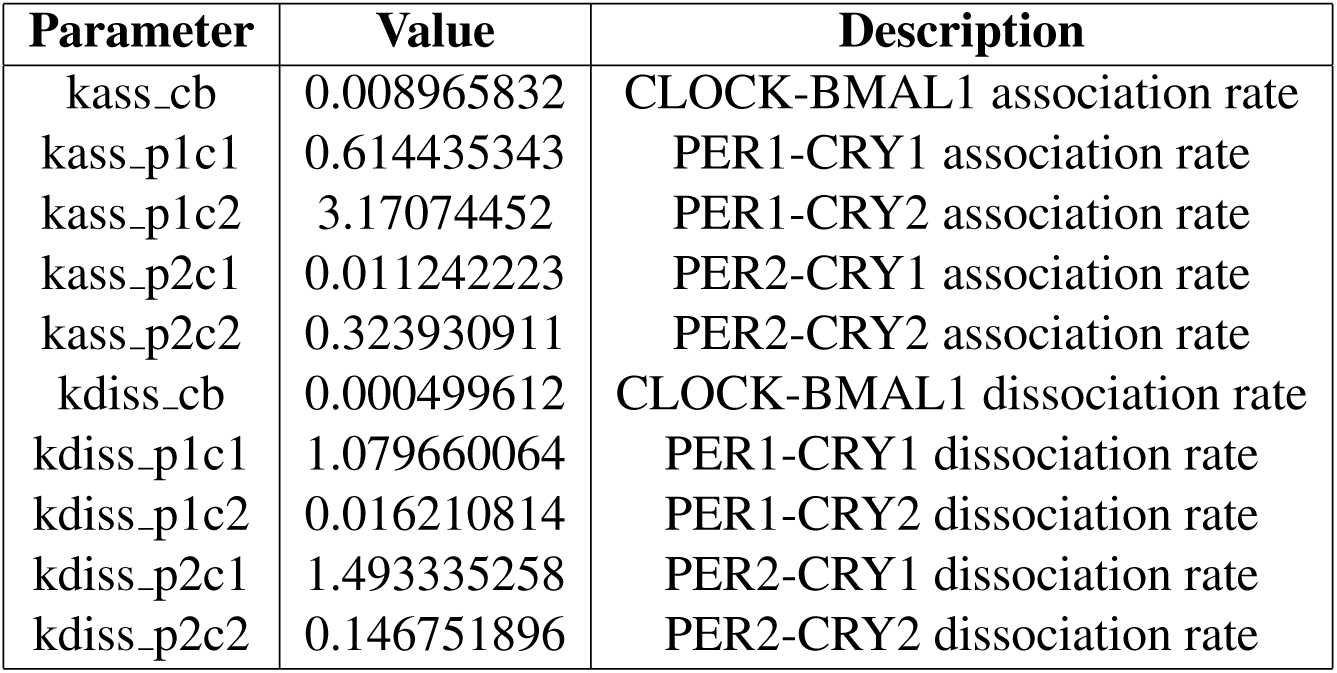
Association and dissociation rates (in hour*^−^*^1^) for the protein complexes.

**Table A6:** Translation rates (in hour*^−^*^1^).

| Parameter | Value | Description |
| --- | --- | --- |
| kp_BMAL1 | 3.271143875 | Bmal1 translation rate |
| kp_CRY1 | 0.738295688 | Cry1 translation rate |
| kp_CRY2 | 0.829860113 | Cry2 translation rate |
| kp_PER1 | 3.798692975 | Per1 translation rate |
| kp_PER2 | 1.980575626 | Per2 translation rate |
| kp_REV | 0.206633966 | Rev-Erb translation rate |
| kp_ROR | 1.875730412 | Ror translation rate |

**Table A7:** Maximal transcription rates (in hour*^−^*^1^).

| Parameter | Value | Description |
| --- | --- | --- |
| V_Bmal1 | 0.646614483 | Bmal1 maximal transcription rates |
| V_Cry1 | 0.401023403 | Cry1 maximal transcription rates |
| V_Cry2 | 0.65185826 | Cry2 maximal transcription rates |
| V_Per1 | 1.109303016 | Per1 maximal transcription rates |
| V_Per2 | 1.671549554 | Per2 maximal transcription rates |
| V_Rev | 0.006423385 | Rev-Erb maximal transcription rates |
| V_Ror | 0.373099089 | Ror maximal transcription rates |
| V_NHE3 | 2.0096808453 | NHE3 maximal transcription rates |
| V_NKCC2 | 0.521355199 | NKCC2 maximal transcription rates |
| V_SGLT1 | 1.0020169907 | SGLT1 maximal transcription rates |
| V_NCC | 0.36248470731 | NCC maximal transcription rates |
| V_ENaC | 0.52277652371 | ENaC maximal transcription rates |

